# *In silico* design of stable single-domain antibodies with high affinity

**DOI:** 10.1101/2024.04.22.589762

**Authors:** Zhongyao Zhang, Rob van der Kant, Iva Marković, David Vizarraga, Teresa Garcia, Katerina Maragkou, Javier Delgado Blanco, Damiano Cianferoni, Gabriele Orlando, Gabriel Cia, Nick Geukens, Carlo Carolis, Alexander N. Volkov, Savvas N. Savvides, Maarten Dewilde, Joost Schymkowitz, Luis Serrano Pubul, Frederic Rousseau

## Abstract

Antibody-based therapeutics have become indispensable in modern medicine, but traditional methods of antibody discovery often present with limitations in developability, cross-reactivity, and ethical concerns. While deep learning and generative approaches have shown promise in the design of high affinity protein binders, *de novo* antibody design remains challenging. Here, we present EvolveX, a structure-based computational pipeline for designing antibody fragments. EvolveX utilizes ModelX and empirical force field FoldX to optimize complementarity-determining regions (CDRs) and TANGO for aggregation analysis. We demonstrate the ability of EvolveX to redesign a single-domain VHH antibody fragment targeting mouse Vsig4 to address two challenges: enhancing stability and affinity for the original target and redesigning it for high affinity to the human ortholog. The redesigned variants of VHH fragments specific to mouse Vsig4 showed improved physicochemical properties, while retaining binding affinities comparable to the original version. Notably, EvolveX improved the binding affinity of VHHs to human Vsig4 by over 1000-fold, transforming low-affinity binders into nanomolar-affinity molecules. Structural analyses by X-ray crystallography and NMR confirmed the accuracy of the designs, which display optimized interactions with the antigen. NGS and re-modelling analysis further demonstrated the efficiency of FoldX-based design pipeline. Collectively, our study highlights EvolveX’s potential to overcome current limitations in antibody design, offering a powerful tool for the development of next-generation therapeutics with enhanced specificity, stability, and efficacy.

## Main Text

Antibodies have become highly promising and dominating tools in both therapeutics and diagnostics. Over the past decades, most of biologics approved by the FDA are therapeutic antibodies ^1^. Traditional antibody discovery methods primarily rely on animal immunization. While effective, these methods present challenges such as suboptimal developability, limited and unpredictable cross-reactivity, and difficulties in targeting conserved binding epitopes ^2,3^. Additionally, ethical concerns regarding the use of animal models further complicate the immunization-based approach ^4^. These limitations have led to a growing shift toward computational-based antibody design and discovery approaches, where various parameters of antibody development can be optimized before experimental testing.

Recent advancements in deep learning networks and diffusion-based generative approaches for designing antibody sequences have attracted increasing attention. However, *de novo* design of high affinity antibodies remains largely unexplored and challenging ^5–8^. In this context, structure-based antibody design has emerged as a promising approach^3,9–13^. Utilizing the computational methods, three-dimensional structure of antibodies and their target antigens can be predicted and modeled, allowing for the further optimization of the full complex structures and their binding interfaces. ^11,12^. Through a combination of computational molecular modeling, rational design of sequences, and experimental validation, structure-based *in silico* antibody design holds the potential to streamline the antibody development process and to facilitate the discovery of antibodies with multiple properties such as enhanced binding affinity and stability ^12,13^.

Here we introduce our innovative *in silico* antibody design pipeline, EvolveX, where the structure optimization is achieved using ModelX ^14–16^, interaction energy and antibody stability are evaluated by the empirical forcefield FoldX ^16,17^ and aggregation propensity is analyzed by TANGO ^18–20^. EvolveX enables the design of antibody sequences that effectively bind to the predefined epitope while maintaining good thermodynamic stability and avoiding the introduction of aggregation propensity. To validate our approach, we employ a process termed *re-paratoping*, where sequence information from a known antibody-protein complex was removed, and new sequences were designed to target the same epitope. The entire pipeline is evaluated through a series of experimental assays, including phage display, biophysical characterization, NMR, X-ray crystallography, and next-generation sequencing (NGS), demonstrating the successful design of stable VHHs with high affinity starting from both high and low-affinity protein complexes ^21^. Overall, our finding underscores the potential of structure-based *in silico* antibody design as a powerful tool in biotechnology and drug discovery.

### *In silico* rebuilding of an antibody paratope starting from a high affinity complex

Wen et al. ^21^ generated a VHH, Nb119 (in this work further referred to as VHH_WT), through classical immunization against the extracellular IgV domain of the mouse Vsig4 protein. This VHH showed high affinity towards mouse Vsig4 (mVsig4, KD ∼5 nM), but a thousand-fold lower affinity towards human Vsig4 (hVsig4, KD ∼5 µM). The structure of the antibody-antigen complex was determined by X-ray crystallography for both orthologs. We used the crystal structure of the complex of the VHH to mVsig4 (VHH_WT-mVsig4, PDBID: 5IMM and 5IMO, ^21^) to validate the capacity of EvolveX to rebuild an antibody paratope starting from a high affinity complex (Fig. 1a). To this end, we removed the sequence information from CDR2 and CDR3 of the VHH_WT- mVsig4 complexes since these regions are most involved in binding to mVsig4: Each amino acid residue in CDR2 and CDR3 (except Proline and Glycine) that resided within 8 Å from the ligand was mutated to alanine. From this all-alanine starting point, the compatibility of each of the selected positions with all 20 amino acids was explored using the *PSSM* function in FoldX, which yields the impact of each point mutation on both the thermodynamic stability of the VHH, as well as interaction energy with the ligand (see methods for details). Based on these numbers, random starting sequences were generated to be compatible with the complex structure. These starting sequences were used as input for a genetic algorithm, where selection was based on the interaction energy computed by FoldX, with preset ceilings for the thermodynamic stability of the VHH (see Fig. 1a and methods for details).

**Fig. 1.**
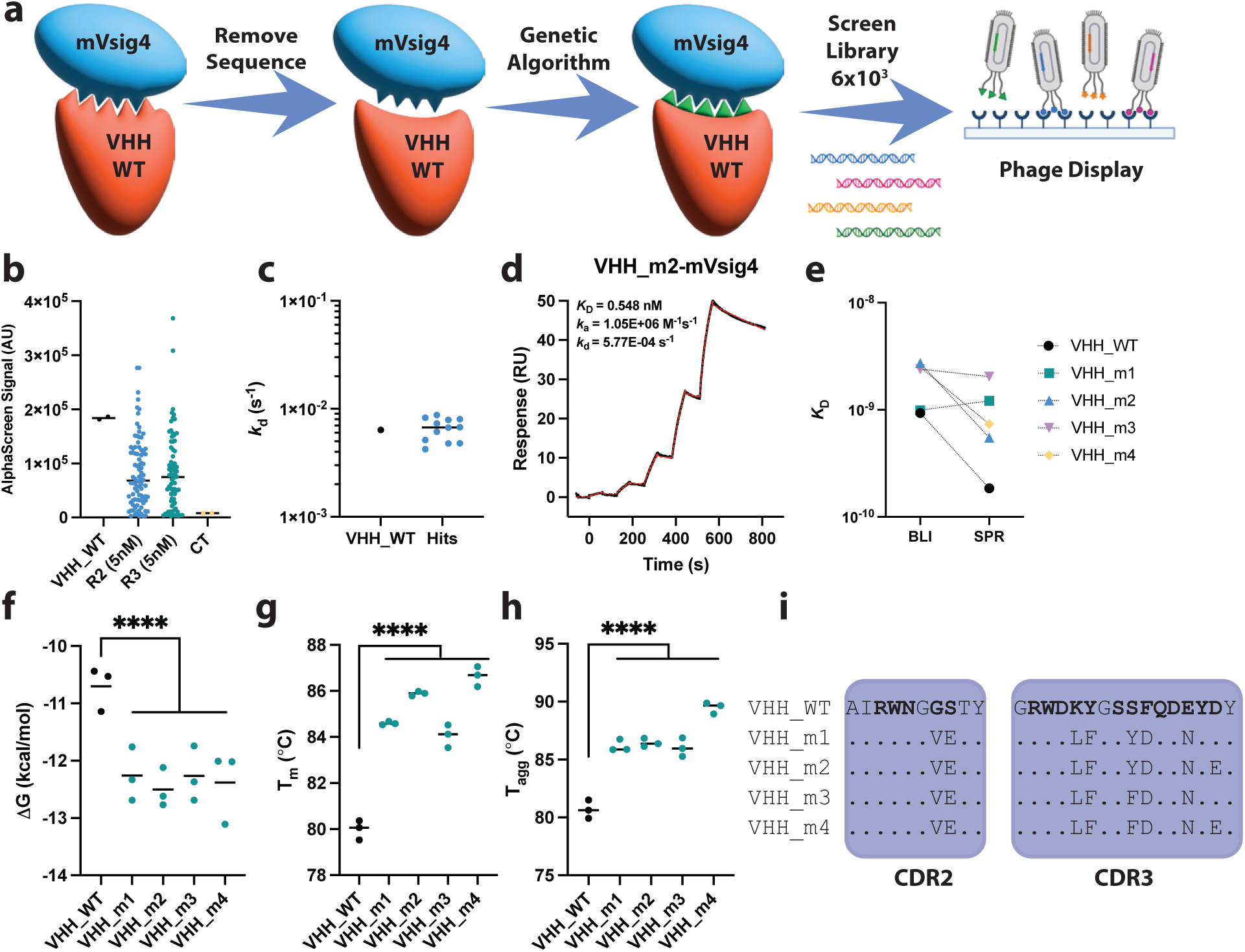
Library design, screening, hit selection and validation of VHH’s designed with EvolveX to bind mouse Vsig4. (**a**) Graphical representation of the EvolveX *in silico* pipeline, where crystal structures of the high affinity complex between mouse Vsig4 (mVsig4) and VHH_WT were used as templates to *in silico* design sequences of CDR2 and CDR3 that should bind mVsig4 with high affinity, while displaying good stability. A small library was subsequently screened using phage display. Icons of DNA strands and phage display were created with BioRender.com. (**b**) Colonies picked after phage display selection were tested for binding to mVsig4 using AlphaScreen. VHH_WT was used as a positive control, colonies picked after two different rounds of selection were screened. (**c**) Off-rates of the top 12 unique sequences based on AlphaScreen signal to mVsig4, determined using BLI. (**d**) Example SPR binding curves of 1 out of 4 hits binding to mVsig4 with low nanomolar affinity. (**e**) Affinity constants of all 4 hits and VHH_WT binding to mVsig4, determined using BLI and SPR. (**f**) ýG values determined through curve fitting based on data shown in G. (**g**) Melting temperatures (Tm) for 4 hits and VHH_WT based on Intrinsic Tryptophan Fluorescence (ITF) through a temperature ramp. (**h**) Aggregation onset temperatures (Tagg) for 4 hits and VHH_WT based on Right Angle Light Scattering (RALS) through a temperature ramp. One way ANOVA was used in (F-H) (ns p>0.123, *p<0.033, **p<0.002, ***p<0.0002, ****p<0.00001). (**i**) Sequence alignment indicating the differences in sequence compared to VHH_WT. CDR2 and CDR3 are indicated by blue boxes, positions identical to VHH_WT are indicated with a dot (.), positions explored with the EvolveX pipeline are displayed bold on the sequence of VHH_WT. AbM CDR definition was used.

The resulting sequences were filtered based on thresholds for the FoldX prediction of interaction energy and VHH stability ^17^, as well as TANGO predicted aggregation propensity ^18^. Additionally, Z-scores were calculated based on densities derived from publicly available crystal structures of VHHs bound to protein ligands, for cavities within the VHH, cavities at the interface of VHHs with their ligand and VHH net charge (see methods for details). The top 6000 sequences with the lowest summed Z-scores were obtained as oligonucleotides and cloned into the VHH_WT backbone in a phagemid vector. The resulting library was enriched for mVsig4 binders through several rounds of phage display using progressively lower concentrations of mVsig4 down to 5 nM (see Supplementary Fig.1 and methods for details). Periplasmic extracts were prepared for 188 randomly picked VHH clones from the enriched libraries, yielding 36 unique sequences (Supplementary Table.1), and subsequently analyzed for mVsig4 binding using the Amplified Luminescent Proximity Homogenous Assay (AlphaScreen) (Fig. 1b). This revealed clones with AlphaScreen signal to noise ratios (S/N ratio) similar- or higher than VHH_WT, of which 12 were selected. Of these VHH variants, we determined the dissociation velocity rate constant governing dissociation from mVsig4 (*k*d) using BioLayer Interferometry (BLI) (Fig. 1c, Supplementary Table.2), which was quite similar in all cases.

The top four candidates (VHH_m1, VHH_m2, VHH_m3, and VHH_m4) with slightly more favorable *k*d values than the rest were subsequently recombinantly produced at larger scale (10 mg), chromatographically purified, and biophysically characterized. Binding affinities (*K*D) for mVsig4 were determined using Surface Plasmon Resonance (SPR) (Fig. 1d and e, Supplementary Fig.3) and BLI (Fig. 1e, Supplementary Fig.2), revealing that the four VHH candidates have affinities close to VHH_WT (Fig. 1e). The four VHHs were subjected to chemical denaturation analysis using urea and guanidine hydrochloride (Supplementary Fig.4). Curve fitting indicated a significant increase in stability by about -2 kcal/mol for all four designed sequences (Fig. 1f) compared to VHH_WT. Thermal protein unfolding (Tm) and aggregation onset temperatures (Tagg) were determined simultaneously using Intrinsic Tryptophan Fluorescence (ITF) (Fig. 1g, Supplementary Fig.5) and Right-Angle Light Scattering (RALS), respectively (Fig. 1h, Supplementary Fig.6). The designed sequences showed increased thermal stability, about 5 °C in both Tm and Tagg, and the relative order in stability coincides quite well between the chemical and thermodynamic denaturation with VHH_m4 being the more stable and VHH_m3 the least.

Although EvolveX computationally explored 5 positions in CDR2 and 12 in CDR3 (∼1×10^22^ theoretical combinations), the four selected candidates were very similar to each other in sequence, differing only at two positions where we found chemically similar amino acids: a Tyr or Phe at position 107 and an Asp or a Glu at position 112, respectively (Fig. 1i, Supplementary Table.3). Interestingly the two more stable VHH sequences VHH_m2 and VHH_m4 contain a Glu at position 112, while having a Phe or a Tyr at position 107 does not seem to have an important role in stability. Compared to wild type, the designed variants carried two mutations in the CDR2 loop and 5 or 6 mutations in the CDR3 loop (Fig. 1i).

The successful identification of the multi-parametric optimized sequences with EvolveX demonstrates the capability of our computational method in designing VHH CDR sequence combinations when starting from a structure of a high affinity complex.

### *In silico* design of optimal CDR2/3 sequences targeting human Vsig4

Subsequently, we set out to identify high-affinity binders starting from a low-affinity complex, starting from structural templates obtained from the crystal structures of the low micromolar affinity complex between VHH_WT and human Vsig4 (PDB ID 5IMK and 5IML, ^21^) (Fig. 2a). The backbone conformation of CDR3 was adapted using the *Bridging* command in ModelX ^16^ using 5IMK as a starting point, adding 14 starting complex structures (see methods for details). Sequence design as described above yielded 6000 candidate sequences (Fig. 2a). The mouse VHH_m and human VHH_h sequences were combined into one phage library, which was selected against hVsig4 using similar procedures as before (Supplementary Fig.1). Based on the enrichment, the phages from the third round of selection with 50nM hVsig4 (R3) and round 4 selection with 5nM hVsig4 (R4) were chosen for screening. We randomly picked 188 colonies from both libraries for expression and sequencing, yielding 112 unique sequences, which we ranked for binding to hVsig4 using classical ELISA. The average ELISA signal from the R4_hVsig4_5nM group is higher than that of the R3_hVsig4_50nM group and most of the clones have a higher ELISA signal than VHH_WT (Fig. 2b, Supplementary Table.4). 56 sequences were tested for off-rate using BLI (Fig. 2c, Supplementary Table.5). The 3 clones with the best off-rates (VHH_h1, VHH_h2 and VHH_h3), the clone with the highest ELISA signal (VHH_h4), and the most occurring clone (VHH_h5) were selected for purification and subsequent biophysical characterization.

**Fig. 2.**
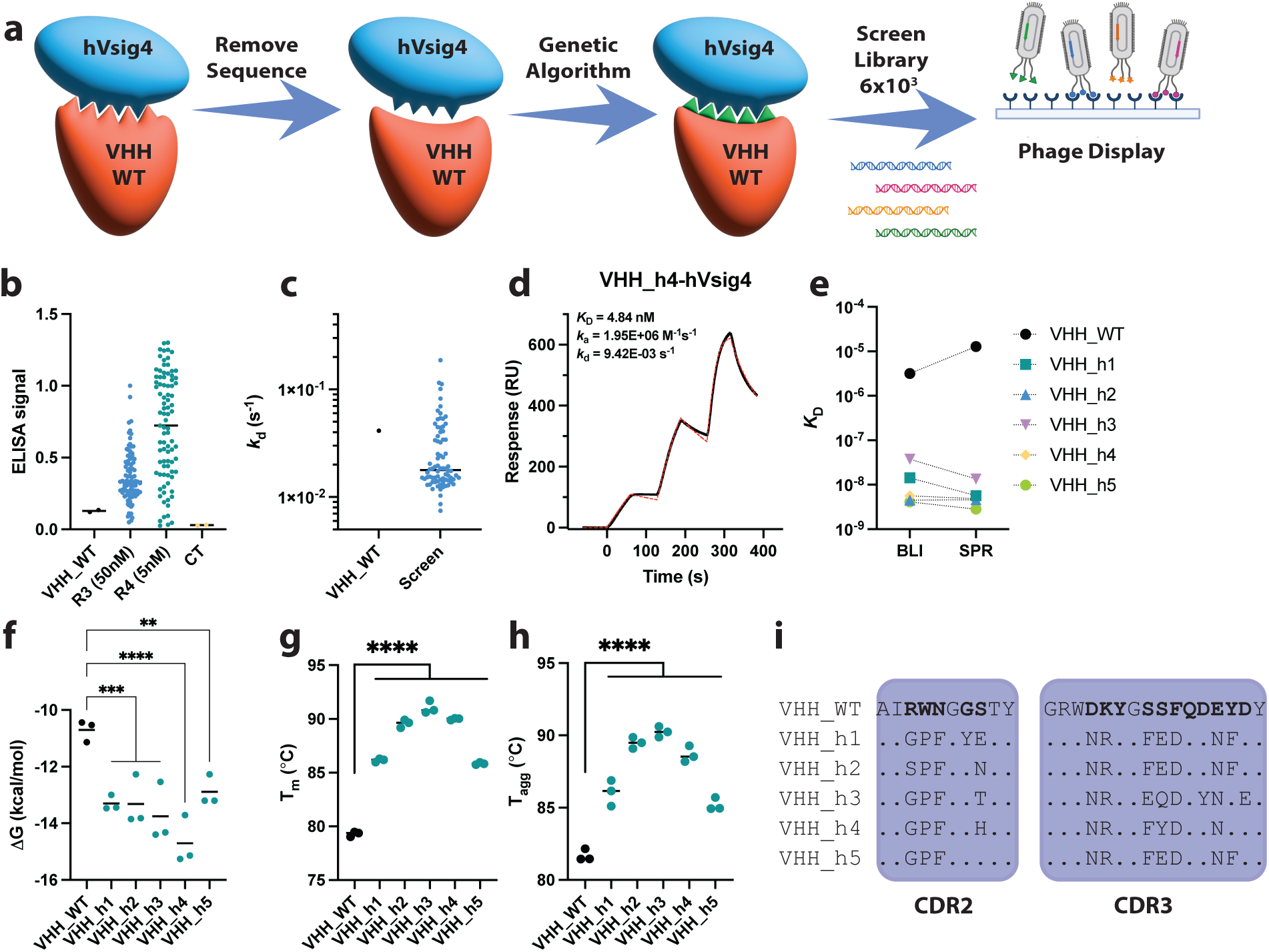
Library design, screening, hit selection and validation of VHH’s designed with EvolveX to bind human Vsig4. (**a**) Graphical representation of the EvolveX *in silico* pipeline, where crystal structures of the low affinity complex between human Vsig4 (hVsig4) and VHH_WT were used as templates to *in silico* design sequences of CDR2 and CDR3 that should bind hVsig4 with high affinity, while displaying good stability. A small library was subsequently screened using phage display. Icons of DNA strands and phage display were created with BioRender.com. (**b**) Colonies picked after phage display selection were tested for binding to hVsig4 using ELISA. VHH_WT was used as a positive control, colonies picked after two different rounds of selection were screened. (**c**) Off-rates of the 82 unique sequences, determined using BLI. (**d**) Example SPR binding curve of 1 out of 5 hits (VHH_h4) binding to hVsig4 with low nanomolar affinity. (**e**) Affinity constants of all 5 hits and VHH_WT binding to hVsig4 determined using BLI and SPR. (**f**) ýG values determined through curve fitting based on data shown in G. (**g**) Melting temperatures (Tm) for 5 hits and VHH_WT based on Intrinsic Tryptophan Fluorescence (ITF) through a temperature ramp. (**h**) Aggregation onset temperatures (Tagg) for 5 hits and VHH_WT based on Right Angle Light Scattering (RALS) through a temperature ramp. One way ANOVA was used in (F-H) (ns p>0.123, *p<0.033, **p<0.002, ***p<0.0002, ****p<0.00001). (**i**) Sequence alignment indicating the differences in sequence compared to VHH_WT. CDR2 and CDR3 are indicated by blue boxes, positions identical to VHH_WT are indicated with a dot (.), positions explored with the EvolveX pipeline are displayed bold on the sequence of VHH_WT. AbM CDR definition was used.

Remarkably, the measured affinity of the designed sequences situated in the low nanomolar affinities (Fig. 2d and e), meaning an improvement in affinity to human Vsig4 of up to 1000-fold over VHH_WT (Fig. 2d and e, Supplementary Fig.7, Supplementary Fig.8). Affinities were cross validated with the VHH_m hits for binding to both mVsig4 and hVsig4. This revealed all designed VHHs show specificity for their intended target (Supplementary Fig.9). All VHHs were subjected to chemical and thermal denaturation. Chemical denaturation by urea (Supplementary Fig.10a) and guanidine hydrochloride (Supplementary Fig.10b) indicated the five designed sequences had an increase in stability from -1.5 up to -3.5 kcal/mol compared with the original human VHH_WT (Fig. 2f and Supplementary Table.6). Both Tm (Fig. 2g, Supplementary Fig.11) and Tagg (Fig. 2h, Supplementary Fig.12) were significantly improved for all designed sequences compared to human VHH_WT (Supplementary Table.6).

VHH_h2, VHH_h4 and VHH_h5 exhibited the strongest binding to hVsig4 (*K*D_BLI and *K*D_SPR ∼ 5 nM), which is 1000-fold better than VHH_WT (*K*D_BLI ∼ 5 µM) (Fig. 2e, Supplementary Table.6). VHH_h1 and VHH_h3 exhibited worse *K*D values (10∼40nM) to hVsig4 than the first three candidates, while still being substantially better than VHH_WT. In all cases we found common mutations among the five VHHs (WN to PF in CDR2, DK to NR, F to D and E to N in CDR3) (Fig. 2i). Interestingly, when comparing the sequences analyzed for hVsig4 and mVsig4, we found some positions that were mutated (although to different residues) in both cases at CDR2 (GS) and CDR3 (D and E). In the case of the hVsig4 we found more positions that were mutated indicating the need of larger sequence changes to ensure high affinity binding.

To confirm the epitope and evaluate the predictions done by EvolveX, the designed VHHs were crystallized in complex with hVsig4 (Fig. 3, Supplementary Fig.13, Supplementary Table.7). All five high-resolution crystal structures revealed that the binding epitope is almost identical among all hit VHHs compared to the crystal structure of VHH_WT (Fig. 3b and c). When superimposing on hVsig4 we see that the orientation for each VHH as well as for VHH_WT is slightly different (Fig. 3b and Supplementary Fig.14). This might be due to the interaction between the designed negative charges in CDR3 (Asp107 and Asp109) and the N-terminus of hVsig4 (Supplementary Fig.14d). Overall, more residues are interacting with hVsig4 in the designed VHHs compared to VHH_WT (Supplementary Table.8). Additionally, the conformation of CDR3 is very similar to the VHH_WT crystal structures (mean pairwise backbone RMSD = 0.29 Å) (Fig. 3c).

**Fig. 3.**
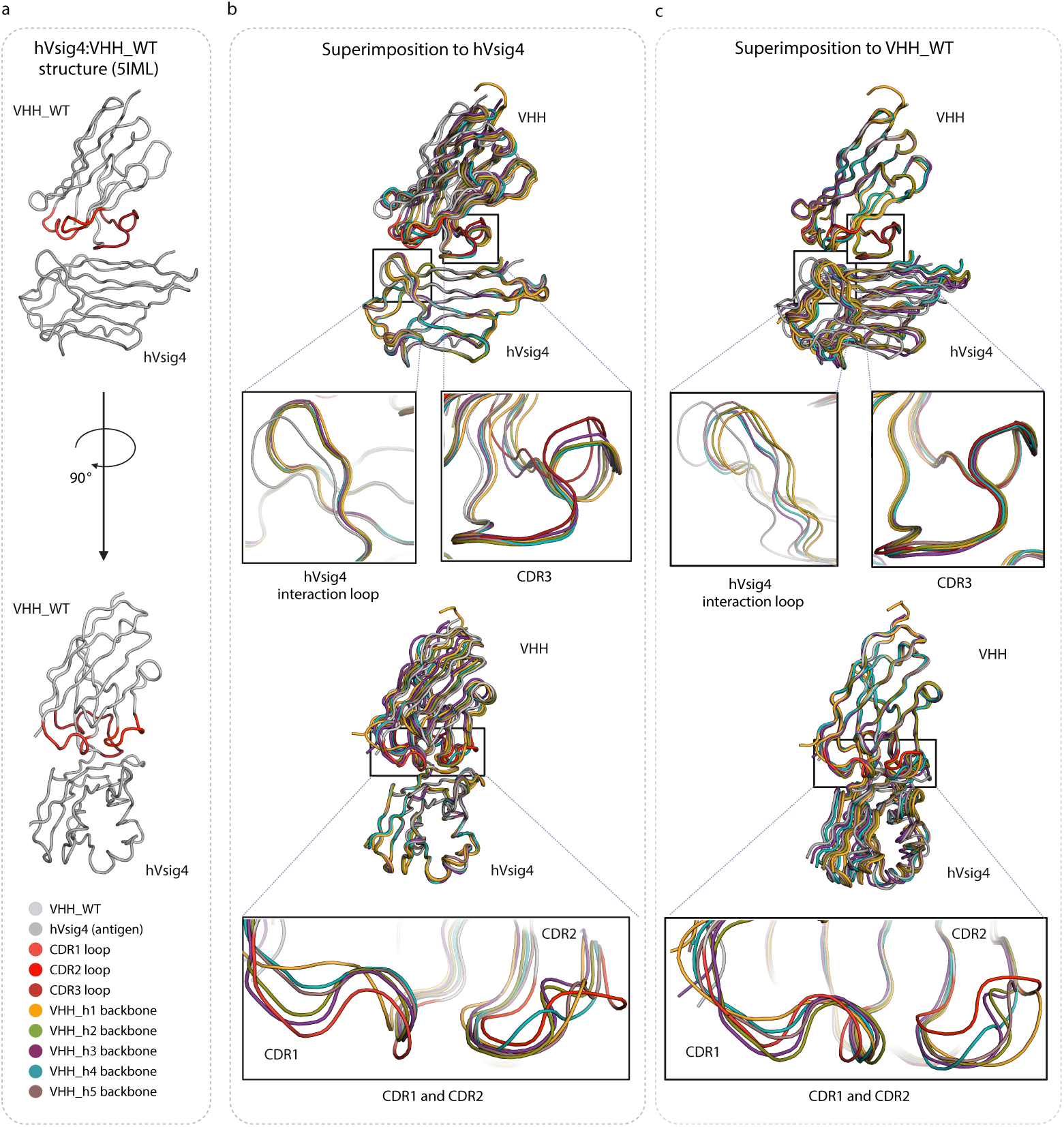
X-ray crystallography high resolution structures of VHH_h1-h5 show epitope retention and conservation of CDR conformation. (**a**) hVsig4:VHH_WT reference structure (PDB:5IML) represented in 2 views with CDRs shown in red. (**b**) Superimposition of the hVsig4:VHH_WT structure with experimental structures of five in silico designed VHHs in complex with hVsig4 aligned to hVsig4 structure. (**c**) Superimposition of the hVsig4:VHH_WT structure with experimental structures of five in silico designed VHHs in complex with hVsig4 aligned to VHH_WT structure. Superimpositions were generated in Pymol using the align function.

To elucidate the biophysical parameters responsible for the increased stability of the designed sequences over the original VHH, we studied VHH_WT and VHH_h4 (the most stable binder to hVsig4) by nuclear magnetic resonance (NMR) spectroscopy. VHH_WT features a well-dispersed heteronuclear single-quantum correlation (HSQC) spectrum, where nearly all peaks could be unambiguously assigned (Supplementary Fig.15). Interestingly, most residues in the three CDRs showed no backbone amide resonances (Supplementary Fig.16). The paucity of the CDR signals was reported before for several other VHHs and attributed to a pronounced flexibility and conformational heterogeneity of the CDR loops ^22–24^. The HSQC spectrum of VHH_h4 is similar to that of VHH_WT, but contains more peaks (Supplementary Fig.17), some of which could be assigned to the residues in the CDR3 loop (Supplementary Fig.18). In addition, several VHH_h4 resonances not observed in the NMR spectra of the VHH_WT (and showing very weak signals in triple-resonance NMR experiments, precluding their unambiguous assignment) could belong to other CDRs. Overall, it appears that VHH_h4 undergoes a much less extensive conformational exchange upon target binding than the WT. Predicted from the backbone chemical shifts, the secondary structures of VHH_WT and VHH_h4 are very similar and agree well with that seen in the X-ray structure (Supplementary Fig.16 and 18), which testifies to the high structural conservation of the VHH fold outside the CDRs. To delve further into differences between VHH_WT and VHH_h4, we performed NMR relaxation analysis (Supplementary Fig.19-21), which revealed that free VHH_h4 in solution maintains the bound-like protein conformation, with closely packed CDR loops undergoing limited motion. At the same time, the CDRs of VHH_WT are largely disordered and involved in extensive conformational exchange. These differences could explain the observed disparity in the thermal stability of the two proteins, where motional disorder of CDRs renders VHH_WT less stable.

### FoldX modeling of VHH hit sequences reveals energy differences between VHH_WT and VHH candidates

After obtaining the sequences and the crystal structures of those Vsig4 candidates, we re-built the structure models of all VHH candidates based on nine available structures (PDBID: 5IMM and 5IMO for VHH_mVsig4, and 5IMK, 5IML, 9EZU, 9EZV, 9EVW, 9EZH, and 9EZI for VHH_hVsig4) using the FoldX *BuildModel* function. The interaction energy (ΔGinteraction) and VHH individual energy (ΔGVHH) were calculated using FoldX *AnalyseComplex* function and visualized in corresponding plots (Supplementary Fig.22). Triplicate running was performed for each model. For mVsig4 hits, VHH_m1 and VHH_m3 exhibited slightly lower ΔGinteraction than VHH_m2 and VHH_m4, consistent with the affinity data from SPR (Supplementary Fig.22a, Fig. 1e). All four hits demonstrated lower ΔGinteraction than VHH_WT, although the difference was not statistically significant ( Supplementary Fig.22a). Additionally, all four hits showed significantly lower ΔGVHH compared to VHH_WT, aligning with the ΔG values measured in the chemical denaturation experiment and Tm/Tagg values measured in the thermal stability experiment (Supplementary Fig.22b, Fig. 1f-h).

For the hVsig4 hits, all five hits showed significantly lower ΔGinteraction and ΔGVHH relative to VHH_WT across all five newly obtained crystal structures (Supplementary Fig.22 c-d). The hit models generated on two original structures (5IMK and 5IML) did not show clear trend relative to VHH_WT (Supplementary Fig.22 c-d). This shows the importance of performing backbone moving to generate the optimized starting structure model when designing from a low-affinity complex.

### FoldX modeling of NGS-identified CDR designs reveals energy change differences between enriched and negative CDR designs

To further investigate the relationship between enriched EvolveX-designed sequences and FoldX-based energy, next-generation sequencing (NGS) was performed on libraries from all rounds of selection (Supplementary Fig.1), including the empty control conducted in parallel with each round (Fig. 4a). As CDR recombination can occur during the PCR steps of the sample preparation, the sequences of CDR2 and CDR3 were extracted and analyzed separately. Enriched CDR designs were filtered from R1 selection for both mVsig4 and hVsig4 based on specific criteria (count > 10, enrichment factor > 10). Notably, the counts for all enriched CDR designs exceeded 1000 (red dots in Fig. 4b, Supplementary Fig.23a, c, i), significantly surpassing the counts for enriched non- designs (black dots in Fig. 4b, Supplementary Fig.23a, c, i), which arose from mutations introduced during cloning or phage selection. An equivalent number of negative CDR designs were identified, defined as sequences present in the input but absent or minimally represented in the R1 selection (Fig. 4c, Supplementary Fig. 23b, d, j). Models incorporating enriched or negative CDR sequences were generated using the FoldX *BuildModel* function based on nine available structures: 5IMM and 5IMO for VHH_mVsig4, and 5IMK, 5IML, 9EZU, 9EZV, 9EZW, 9EZH, and 9EZI for VHH_hVsig4. The interaction energy change (ΔΔGinteraction) and VHH individual energy change (ΔΔGVHH) were calculated using FoldX *AnalyseComplex* function and visualized in corresponding plots (Fig. 4d-e, Supplementary Fig.23e-h, k-l).

**Fig. 4.**
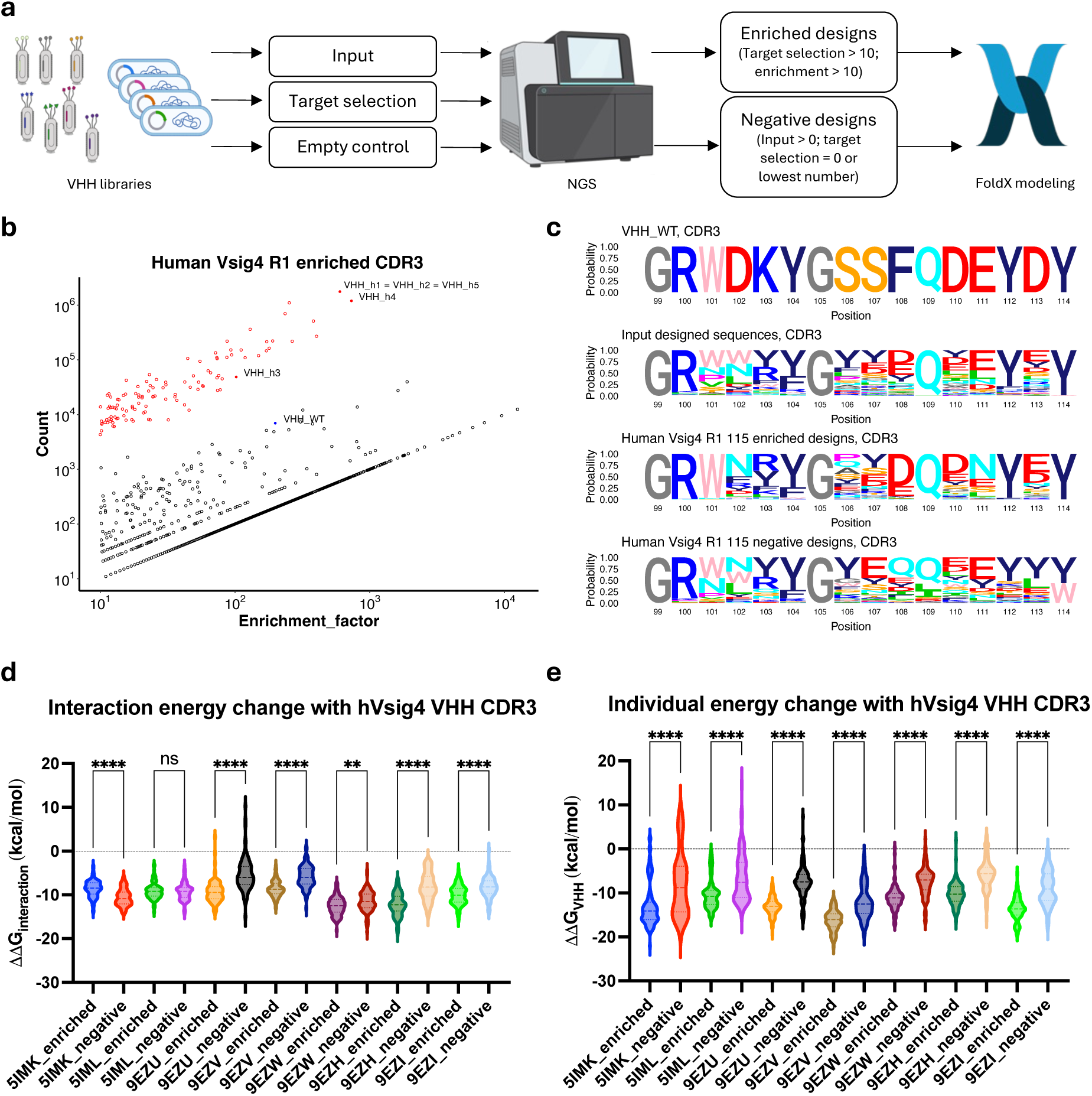
FoldX modeling of NGS-identified hVsig4 CDR3 designs reveals energy change differences between enriched and negative CDR designs. (**a**) Graphical representation of the NGS preparation, analysis and FoldX modeling workflow. Input, target selection, and empty control libraries were analyzed by NGS. Enriched and negative CDR designs were filtered based on specific criteria and subsequently modeled using the FoldX force field. Icons of phages and NGS sequencer were created with BioRender.com. (**b**) Count and enrichment factor of human Vsig4 enriched CDR3 sequences (count > 10, enrichment factor > 10, enrichment factor is the ratio of the count in target selection to the count in empty control). Red dots are the sequences designed by EvolveX, red solid dots are the sequences from identified hits, blue solid dot is VHH_WT, and black dots are non-designs or mutants generated during the PCR or phage selection stages. (**c**) Sequence logo plots of VHH_WT CDR3, input CDR3 designed sequences, 115 hVsig4 enriched CDR3 designed sequences and 115 negative CDR3 designed sequences. AbM CDR definition was used. (**d**) Comparison of the interaction energy change (ΔΔGinteraction) between enriched and negative hVsig4 CDR3 designs. One way ANOVA was run between each group (ns p>0.123, *p<0.033, **p<0.002, ***p<0.0002, ****p<0.00001). (**e**) Comparison of the VHH individual energy change (ΔΔGVHH) between enriched and negative hVsig4 CDR3 designs. One way ANOVA was run between each group (ns p>0.123, *p<0.033, **p<0.002, ***p<0.0002, ****p<0.00001).

For mVsig4, 31 enriched CDR2 designs and 34 enriched CDR3 designs were identified (Supplementary Fig.23a-d). Compared to negative CDR sequences, all models with enriched CDR2 designs showed significantly lower ΔΔGinteraction and all models with enriched CDR3 designs showed significantly lower ΔΔGVHH on both VHH-mVsig4 structures (Supplementary Fig.23e-h). For hVsig4, 38 enriched CDR2 and 115 enriched CDR3 designed sequences were identified and analyzed across all seven VHH_hVsig4 structures (Fig. 4b-e, Supplementary Fig.23i-l). While no significant differences were observed between enriched and negative CDR2 models in ΔΔGinteraction and ΔΔGVHH, the average ΔΔGinteraction across all enriched CDR2 designs was consistently lower than that of the negative models (Supplementary Fig.23k-l). In five newly obtained structures (9EZU, 9EZV, 9EZW, 9EZH, and 9EZI), enriched CDR3 models exhibited significantly lower ΔΔGinteraction (Fig. 4d). In addition, all enriched CDR3 models demonstrated significantly lower ΔΔGVHH compared to the negative models across all analyzed structures (Fig. 4e). Overall, the results demonstrated that FoldX parameters were highly effective in distinguishing optimal designs from negative sequences on the proper structures. Beyond the top hits we identified, 30–100 optimal designs were successfully obtained for each CDR based on both NGS and FoldX analysis.

## Discussion

The advent of computational methods for antibody design is revolutionizing the field, allowing for the targeted creation of antibodies with specific binding capabilities and stability. In this study, we demonstrate the efficacy of an *in silico* antibody design pipeline, EvolveX, in designing VHHs targeting predefined epitopes on protein targets. Our results highlight the versatility and robustness of the approach, as evidenced by successful sequence design efforts starting from both high and low-affinity complexes.

The success of the *in silico* antibody design pipeline is underscored by the validation experiments conducted for both high and low-affinity complexes of VHH_WT with mouse and human Vsig4, respectively. Through a combination of sequence and structure optimization, and biophysical and structural experimental validation, EvolveX was able to design small, targeted libraries yielding high-affinity VHHs. Notably, the designed VHHs consistently exhibited significantly enhanced thermal, colloidal, and chemical stability, indicating the robustness of the computational design process. Interestingly, comparison of the designed VHHs targeting mouse and human Vsig4 revealed differences in the mutation patterns within the complementarity-determining regions (CDRs), indicating the need for tailored design strategies depending on the target species.

The designed VHHs exhibited not only high affinity but also demonstrated thermodynamic stability comparable to or better than the wild-type antibody. Structural analysis of the VHH- hVsig4 protein complexes confirmed epitope retention and revealed insights into the binding mechanism, providing valuable information for further optimization and development.

Furthermore, FoldX modeling of VHH hit sequences and NGS-identified sequences demonstrated consistency between wet-lab data and *in silico* predicted parameters. Models based on low-affinity hVsig4-VHH complex (5IMK) showed no clear trends or, in some cases, opposite results to expectations. However, the backbone of VHH_WT in the 5IMK complex was initially adapted using ModelX during the design process to include additional starting structures, which could prove highly valuable in identifying high affinity hVsig4 hits. These findings emphasize the critical importance of backbone conformation exploration into the modeling workflow and highlight the need for further refinement of the force field to enhance predictive confidence.

Collectively, our study demonstrates the power of computational antibody design in generating VHHs with desired binding properties. The combination of molecular modeling, sequence optimization, and experimental validation offers a systematic and efficient approach for antibody engineering, with broad implications for drug development, diagnostics, and biotechnology. Future work will focus on further refining the design and modelling pipeline, reducing library sizes and exploring its application to additional protein targets and therapeutic modalities.

## Acknowledgments

Thanks for support to the Unit of Structural Biology (IBMB-CSIC), to the Xaloc beamline at ALBA and to the Crystallography Platform at the Barcelona Science Park (PCB). We thank the staff of the EMBL beamline P14 (PETRA III, Hamburg, Germany) for beamtime allocation and technical support.

## Methods

### Sequence Design: Template generation

The original PDB structures of the wildtype VHH were downloaded for the PDB. Two low affinity complexes of wildtype VHH bound to human Vsig4 (hVsig4; 5IMK and 5IML) and two high affinity complexes of wildtype VHH bound to mouse Vsig4 (mVsig4; 5IMM and 5IMO). Sidechains were subsequently optimized using the FoldX ^17^ command *RepairPDB*. Backbones were optimized using the ModelX ^14–16^ command *Vibrate.* It generates 1 by 1 Degree rotations in a range for -20 to 20 Degrees around an imaginary axis along N and C for every amino acid, approximating phi, psi torsional rotations but reducing the number of rotations to one angle. The human bound structure 5imk was the starting point to explore alternative backbones for CDR3 using the *Bridging* command in ModelX. ModelX allows to digest customized peptide fragments databases adapted to the different modeling requirements. The peptide fragments were used to graft CDR3 loop conformations generating conformational variability, in our case we created *in situ* a database using 120000 X-ray structures with resolution <= 2.5 Å. 18544 backbone moves were generated and selected using these criteria, first they were filtered out those having backbone-backbone clashes and then selected those having a number of contacts bigger than 36, the remaining 85 models that passed the filters were ranked according to their FoldX interaction energy and 14 were selected, resulting in 14 structures with slight movements in CDR3. The 22 structures were subjected to *de novo* sequence design of CDR2 and CDR3.

### Sequence Design: Explore positions

Each amino acid part of CDR2 or CDR3 (according to Chothia numbering scheme ^25^) and within 8 Ångström from the ligand was explored separately for each possible amino acid using *BuildModel*, after which the effect of each mutation on stability and interaction energy was calculates using the FoldX commands *BuildModel* and *AnalyseComplex*, respectively.

### Sequence Design: Reduce search space

To reduce the sequence space, the positions were filtered. Positions were removed where only the wildtype did not result in major destabilization of the VHH (ddG_binder < 1). Additionally, positions where the wildtype amino acid was best for VHH stability (ddG_binder rank = 1) and where mutations had little effect on interaction energy (ddG_AC variation < 0.1). Positions where the wildtype amino acid is Proline were not considered either. Mutations that destabilize the VHH (ddG_binder > 1 kcal/mol) or reduce the interactions energy (ddG_AC > 1kcal/mol) were removed from the search space.

### Sequence design: Genetic algorithm (EvolveX)

Starting sequences were randomly generated based on the remaining search space. Each starting sequence is evolved through a process based on a genetic algorithm ^26^. The 50 start points (threads) for each template are used as input for the genetic algorithm. The genetic algorithm was run for 500 generations. For each generation 10% of the 50 are recombined with each other and 90% is subjected to a point mutation. Recombination is done by randomizing the identities of each parent for each explored position and splitting the resulting list of mutations at a random point. Point mutations are selected for a random position, while selecting a new amino acid for that position semi-randomly according to a frequency table based on amino acid distributions of paratopes ^27^. The effect of the mutation event is calculated using FoldX commands *BuildModel*, *AnalyseComplex*, *Stability*, TANGO ^18^. Thresholds are set manually for VHH stability (dG_binder) and VHH intraclashes (IntraClash_binder). A mutation in rejected if the difference with the parent is positive and the value of the mutant is above the manually set threshold. When these criteria are met, the main driver is interaction energy (ddG_AC < 0 kcal/mol) which allows the mutant to be accepted and the metropolis criterion is used when the interaction energy is decreased (ddG_AC >= 0 kcal/mol). If the mutant is accepted it is used as input for the next generation, if it is rejected the parent is used as input.

### Sequence selection

R was used to select from the 500K accepted sequences. Only sequences are considered that have a satisfactory interaction energy with the ligand (dG_AC < -17.5 kcal/mol), good VHH stability (dG_binder < -5 kcal/mol), low predicted aggregation propensity (total TANGO score < 450). For the remaining models, VHH intracavities (Cav_binder), interface cavities (Cav_Int) and VHH net charge (ChAb) were calculated using YASARA ^28^. Z-scores were calculated for all three variables based on their distributions in known VHH complexes in the PDB. Models with Z-scores above 2 for any of the three variables were excluded. The three Z-scores were summed. The models with the lowest summed Z-score and designed to bind mouse Vsig4 were selected to be included in the library (AffinityMouse), with the highest summed Z-score were selected as negative control (NegativeControlMouse). The models with the lowest summed Z-score and designed to bind human Vsig4 were selected to be included in the final library (AffinityHuman), 207 with the highest summed Z-score were selected as negative control (NegativeControlHuman).

### Vsig4 preparation

Mouse recombinant Vsig4 (mVsig4) used in phage selection, screening, and VHH affinity measurement was bought from Sino Biological (50187-M08H-100) and biotinylated with EZ-Link NHS-PEG4-Biotin (Thermo Scientific, A39259) according to the manufacturer instructions. The mVsig4 material used for crystallization (amino acids 20-139, with C-terminal His tags) was cloned into plasmid pVDS101 (a modified version of pHEN6), expressed in TG1 *E. coli* strain, and purified from the periplasmic extract by a Ni-charged column followed by a size exclusion chromatography (SEC) (Superdex 75 16/600 column).

Human recombinant Vsig4 (hVsig4) used in phage selection, screening, and VHH affinity measurement (amino acids 20-137, with C-terminal Avitag and His tags) was cloned into pVDS101 plasmid and expressed in AVB101 *E. coli* strain (AVIDITY) to allow *in vivo* biotinylation, following the suggested protocol. Then the protein was extracted from periplasm and purified by Ni-NTA chromatography. The hVsig4 protein used in crystallization (amino acids 20-137, with C-terminal His tags), was cloned into pLTDA104 plasmid (also a modified version of pHEN6), expressed in TG1 *E. coli* strain and purified following the same procedures with mVsig4.

### Cloning of the synthetic Vsig4 CDR23 VHH library

The oligo pool containing 12,405 sequences (12,405 CDR pairs, 3200 unique CDR2s, 9884 unique CDR3s), encoding for the region from CDR2 to CDR3 of the VHHs, flanked by two restriction sites (BpiI and Eco91I) was ordered at Twist Biosciences. The oligo pool was amplified with a KAPA HiFi HotStart ReadyMix PCR Kit (Roche, KK2602). The amplified product was then restriction digested and ligated into the phagemid vector pP002ph008 (a modified version of pMECS, which encodes for the non-library encoded parts of VHH_WT, followed by 3xFlag/6xHis tag at the C-terminus of the VHH insertion site) with T4 DNA ligase (Thermo Scientific, EL0011) at 16°C overnight. The purified ligation sample was transformed into the TG1 electrocompetent cells (Lucigen, 60502-2). The resulting library had a size of >10^7^ CFU and a high insertion rate (> 90 %). At last, the phage library was produced with the infection of M13KO7 helper phage (Thermo Fisher Scientific, 18311019) according to the protocol published by Pardon et al^29^.

### Phage library selection

The VHH displaying phage library was selected against multiple rounds of mVsig4 or hVsig4 according to the standard protocol ^29^ (Supplementary Fig.1 and Supplementary Fig.23). Briefly, 2×10^11^ CFU of phages, biotinylated antigens, and streptavidin magnetic beads (Thermo Scientific, 88817) were firstly blocked at PBS buffer with 2% of milk for at least 0.5h at room temperature. Then the blocked phages and antigens were co-incubated for 2h at room temperature to allow the binding of VHH-displaying phages with the targets. Non-binding phages were discarded with 5 consecutive washing steps with PBS/2% milk, whereas bound phages were captured by the streptavidin beads and eluted by trypsin. The eluted phages were rescued with the exponentially grown TG1 cells for the next round of selection. In the end, the output phage libraries from desired rounds of selection were sub-cloned into the vector pLTDA118 (a modified pHEN6 vector with C-terminal 3xFlag/6xHis tag) and transformed into TG1 cells. Single colonies were randomly picked to grow in the 96-well deep-well plate for the preparation of periplasmic extracts, and Sanger sequenced as previously described ^30^.

### AlphaScreen

AlphaScreen FLAG (M2) detection kit (PerkinElmer, 6760613M) was used for the initial screening of the mVsig4 hits. In short, in a 384-well Optiplate (PerkinElmer, 6007290), 5µL of diluted biotinylated mVsig4, 5µL of diluted periplasmic extracts in a 96-well plate from different clones, and 10µL of anti-FLAG conjugated acceptor beads (50µg/ml) were incubated together for 1h at room temperature in the dark. Then 5µL of streptavidin donor beads (100µg/mL) was added and incubated in the dark for 25-30 minutes. The signals were read (excitation 680nm; emission: 520-620nm) using the EnVision multilabel plate readers (PerkinElmer).

### ELISA screen

Human Vsig4 hits were first screened in an ELISA assay. In brief, 100µL of biotinylated hVsig4 (2µg/mL) was coated in the ELISA plate in advance. The plates were blocked with PBS/1% BSA for 2h at room temperature. Subsequently, 100µL of periplasmic extracts were added per well, and incubated for 1.5h at room temperature. Then 100µL/well of anti-FLAG-HRP antibody (Sigma-Aldrich A8592) (1:2500 dilution) was added and incubated for 1 more hour. At last, TMB solution (50µL/well) was used to detect and after 1-5 minutes, 50µL/well of H2SO4 was added to stop the reaction. The plates were then read at 450nm of wavelength in a plate reader.

### Biolayer interferometry (BLI) off-rate assay

The clones with unique sequences and significant AlphaScreen or ELISA signal (signal/noise, S/N > 3) were further selected, re-arrayed in a 96-well plate, and screened by a biolayer interferometry (BLI) off-rate assay on Octet RED96 (Sartorius). Ni-NTA biosensors (Sartorius, 18-5101) were first used to assess the expression of different clones. After equilibration of the biosensors for at least 10 minutes in 1x Sartorius/Octet kinetic buffer and an extra 120 seconds for the baseline, the biosensors were dipped into 10x diluted periplasmatic extracts for 300 seconds, followed by 300 seconds of dissociation in 1x kinetic buffer. Afterward, streptavidin (SA) biosensors (Sartorius, 18-5019) were used to measure the off rate of different clones. With the same pre-wet procedure and the baseline setting, biotinylated Vsig4 material was first loaded on the biosensors, after which, the tips were sequentially submerged in baseline wells, association wells with diluted periplasm samples (300s), and finally into the dissociation wells (300s). Data was processed on the Octet RED analysis software.

### VHH expression and purification

VHH candidates used for initial affinity measurement were in the vector pLTDA118 (a modified pHEN6 vector with C-terminal 3xFlag/6xHis tag) and expressed in the TG1 cells following the standard protocol ^29^. The purification of these VHHs was done by either the Ni-NTA chromatography or the AmMag™ SA Plus System (Genscript, L01013) with Ni magnetic beads (Genscript, L00776). VHHs used in stability study and crystallization were cloned into the vector pLTDA104 (a modified pHEN6 vector with C-terminal 6xHis tag) and expressed in the TG1 cells following the same procedure. These VHHs were then purified by a Ni-charged column followed by size exclusion chromatography (SEC).

### Biolayer interferometry (BLI) affinity measurement

Purified VHHs were used to determine the affinity with mVsig4 and hVsig4 on Octet RED96 (Sartorius). Briefly, streptavidin (SA) biosensors (Sartorius, 18-5019) were first equilibrated for at least 10 minutes in 1x Sartorius/Octet kinetic buffer. Subsequently, the tips were dipped into the biotinylated Vsig4 (3-5µg/mL) target for 30-60 seconds. Afterward, the tips were sequentially submerged in the 1x kinetic buffer for the baseline (120s), VHH dilution samples for the association (300s), and in the end back to the 1x kinetic buffer for the dissociation (300s). Fitting and binding kinetics determination was performed with a 1:1 model interaction on the Octet RED analysis software.

### Surface plasmon resonance (SPR) affinity measurement

Binding kinetics and affinity of various VHHs for the human and murine Vsig4 were evaluated by surface plasmon resonance on a BIAcore T200 instrument (Cytiva) with a running buffer composed of 10mM HEPES, 150mM NaCl & 0.005% Tween 20. The assay format involved ligand capture on a Sensor S Sensor Chip SA chip. Briefly, the biotinylated ligands were immobilized by non-covalent capture (binding to streptavidin) following the instructions provided with the Cytiva chip. mVSIG4 was captured on flow cell 2, and hVSIG4 was captured on flow cell 3, leaving flow cell 1 as a subtractive reference. The capture level of the mVsig4 was targeted between 100 and 200 resonance units, and the hVsig4 was targeted between 1200 and 1300 resonance units. A serial dilution of the VHHs was flowed over the immobilized Vsig4 (50μL/min for 1 minute) and allowed to dissociate for 4 minutes. The capture surface was regenerated with a 60-s injection of 3 M MgCl2 (50 μL/min for 1 min). A 3-fold concentration dilution series of each VHH variant ranging was used to analyze binding to mVsig4. For the VHH_WT, VHH_m1-VHH_m4 the concentration series ranged from 0.25nM to 20nM. For the VHH_h1-VHH_h5 the concentration dilution series ranged from 6.2nM to 500nM. A 3-fold concentration dilution series of each VHH variant ranging was used to analyze binding to hVsig4. For the WT and VHH_m1, the concentration dilution series ranged from 3.7nM to 300nM. For VHH_h1-VHH_h5 and VHH_m4 the concentration dilution series ranged from 0.25nM 20nM. For the VHH_m2 and VHH_m3, the concentration dilution series ranged from 2.1nM to 167nM. All sensograms were analyzed using a 1:1 Langmuir binding model with software supplied by the manufacturer to calculate the kinetics and binding constants.

### Thermal stability measurement

The temperature at which 1.0 mg/mL VHHs unfold (Tm, unfolding temperature) and aggregate (Tagg, aggregation onset temperature) using an UNcle device (Unchained Labs, CA, USA). More specifically, the Tm and Tagg were determined simultaneously via intrinsic fluorescence (ITF) and static light scattering (SLS) measurements, respectively, during gradual heating from 15 to 95°C at 0.3°C/min. The excitation wavelength of 280 nm was used for both measurements, with the emission wavelength ranging from 250 to 720 nm for the intrinsic fluorescence and corresponding to 266 nm for the SLS. The intrinsic fluorescence values between 320 and 370 were used to calculate the barycentric mean values (BCMs), resulting in a smooth curve. Curve fitting with R- studio was used to calculate the Tm value corresponding to the inflection point of the sigmoid curve. The Tagg value was determined using manually set thresholds, to allow for proper comparison between variants, ignoring baseline noise. All samples were measured at least in triplicate.

### Chemical stability measurement

Chemical denaturation was performed using both Urea (0-9.2M) and Guanidine Hydrochloride (GdnHCl, 0-5.5M) using a gradient of 24 steps. Pipetting the different steps was done using a robot (Opentrons OT-2) from a protein stock solution of 1 mg/mL in PBS with a final concentration of 0.04 mg/mL and freshly prepared denaturant stock solution of 10M Urea or 6M GdnHCl. All stock solutions were filtered using 0.22 μM filter. After pipetting, plates were spun down in a centrifuge at 1000g for 2 minutes. The plates were incubated for 2 hours at 25°C with a lid to avoid evaporation of the sample. Samples were measured using the standard protocol on the SUPR-CM (ProteinStable).

### Crystallization of the VHH:hVsig4 complexes

For VHH_h1, VHH_h2 and VHH_h3 complexes with hVsig4, 1.2 molar excess of VHH was added to hVsig4, incubated for 15 minutes at room temperature and purified by SEC on a Superdex 75 Increase 10/300 (Cytiva) column using HBS pH 7.4 buffer. Fractions containing the complex were pooled and concentrated to 4 - 6 mg/ml. Crystallization screens were set up using a Mosquito liquid handling robot (TTP Labtech) in sitting-drop format using 100 nl of protein mixed with 100 nl of mother liquor in SwissSci 96-well triple drop plates incubated at 287 or 294 K. VHH_h2:hVsig4 complex hit was obtained by micro-seeding using crystal seeds of VHH_h1:hVsig4 complex grown in condition 0,17M ammonium sulfate, 15% glycerol, 30,5% PEG 4000. Seed stock was prepared by crushing the crystals with a loop in original mother liquor drop. Crystal seeds were added using a cat whisker to drops containing 100 nl of protein mixed with 100 nl of mother liquor in multiple conditions. Plate was incubated at 287K. Best diffracting crystal of this complex was obtained in 0,1M Na-citrate pH 5,6, 20% 2-propanol and 21% PEG 4000. Crystals were cryoprotected in mother liquor containing glycerol or ethylene glycol prior to being cryocooled in liquid nitrogen. Diffraction data was collected at 100 K at the P13 microfocus beamline at PETRA III (Hamburg, Germany).

For VHH_h4 and VHH_h5 complexes with hVsig4, VHH and hVsig4 were mixed in a 1:1 molar ratio respectively, incubated for 40 minutes at room temperature and purified by size exclusion chromatography (SEC) on a Superdex 75 Increase 10/300 (Cytiva) column using 20mM Tris pH 7.5, 5% Glycerol and 150mM NaCl buffer. Fractions containing the complex were pooled and concentrated to 3.5-4.5 mg/ml. The crystallization screens were set up using the hanging drop vapor-diffusion technique by mixing 1 µl of protein solution with 1 µl of mother liquor. For this purpose crystallization conditions consisting of 0.1M Bis-Tris pH 6.0-6.5 and 25-30% PEG 3350 were used, based on the previously reported conditions that yielded crystals of native VHH_WT:hVsig4 complexes ^21^. Plates were incubated at 293K. Obtained crystals were cryoprotected in mother liquor containing 15% glycerol prior to being cryocooled in liquid nitrogen. Diffraction data was collected at 100 K at the Xaloc beamline at ALBA (Barcelona, Spain). Detailed crystallography setup and data collection statistics are summarized in Supplementary Table.7.

### Determination of crystallographic structures of the VHH:hVsig4 complexes

Datasets for complexes of VHH_h1, VHH_h2 and VHH_h3 were integrated and scaled using XDS ^31^ and AIMLESS and the data quality was analyses by Phenix Xtriage and datasets for complexes of VHH_h4 and VHH_h5 with hVsig4 were processed with autoPROC ^32^.

Initial phases for the VHH:hVsg4 complexes were obtained by maximum-likelihood molecular replacement with Phaser ^33^ using the coordinates of native structures of hVsig4 and VHH (PDB ID 5IML) as a search model. A model for each of the VHH variants was manually rebuilt in Coot ^34^ and the refinement of coordinates and atomic displacement parameters for all the structures was performed in Refmac ^35^ and Phenix refine ^36^. Model and map validation tools in Coot and the PHENIX suite were used throughout the workflow to guide improvement and validate the quality of crystallographic models. Refinement statistics are summarized in Supplementary Table.7.

### NMR spectroscopy

The NMR samples contained 0.8-1 mM [^13^C,^15^N] or ^15^N VHH in 20 mM sodium phosphate 50 mM NaCl pH 6.5, 0.02% NaN3, and 10 % D2O for the lock. All NMR spectra were acquired at 313 K on a Bruker Avance III HD 800 MHz spectrometer, equipped with a triple-resonance TCI cryoprobe for enhanced sensitivity. Three-dimensional experiments were acquired with a non-uniform sampling (25-50%) as implemented in TopSpin 3.7 (Bruker). The NMR data were processed in TopSpin 3.7 (Bruker) or MddNMR ^37^ and NMRPipe ^38^, and analyzed in CCPNMR^39^. Assignments of protein backbone resonances were obtained from [^13^C,^15^N]-labelled samples of C-terminally triFlag- and His-tagged VHH_WT and VHH_h4 by establishing intra- and inter-residue connectivities in 3D HNCACB, HN(CO)CACB, HNCO, HN(CA)CO, and HBHA(CO)HN spectra at the corresponding peak frequencies in the 2D [^1^H,^15^N]-HSQC spectrum. The assignments were subsequently transferred to [^1^H,^15^N]-HSQC spectra of ^15^N-labelled C-terminally His-tagged VHH_WT and VHH_h4 constructs and verified by 3D ^15^N-NOESY-HSQC (mixing time 120 ms). The assigned chemical shift lists were deposited in Biological Magnetic Resonance Data Bank (BMRB) entries 52361 (VHH_WT) and 52360 (VHH_h4). The VHH secondary structure (Supplementary Fig.16 and Supplementary Fig.18) was predicted from the backbone chemical shifts using the CSI function and the DANGLE module ^40^ in CCPNMR ^39^ or the web-based RCI ^41^.

Relaxation measurements were performed on ^15^N-labelled samples of C-terminally His-tagged VHH_WT and VHH_h4. Longitudinal (R1) and transverse (R2) relaxation rates were obtained from [^1^H,^15^N] HSQC-based pseudo-3D spectra with interscan delays of 2.5 s. The inversion recovery experiment (R1) had relaxation delays of 10, 100, 1000, 400, 600, 800, 500, 1200, 200,50, 100, 300, 900 ms (in this order), while Carr-Purcell-Meiboom-Gill (CMPG) measurments (R2) employed relaxation delays of 17, 68, 136, 170, 85, 34, 51, 34, 119, 153, 102 ms (in this order). Steady-state, heteronuclear {^1^H}-^15^N NOEs were measured in an interleaved experiment with and without ^1^H saturation and the relaxation delay of 6 s and performed in duplicate to estimate the experimental errors. For each analysed [^1^H,^15^N]-HSQC peak, the R1 and R2 values were obtained from exponential decays of signal intensities in CCPNMR ^39^, while {^1^H}-^15^N NOEs were determined as ratios of peak intensities in the spectra with and without ^1^H saturation. Relaxation analysis was performed with Modelfree v 4.20 ^42^ using FastModelfree setup ^43^. Initial estimates of the diffusion tensor parameters were obtained with quadric_diffusion ^44^ from the experimental R2/R1 data that excluded the residues with high R2 values (R2 ≥ <R2> + σR2) as likely exhibiting significant internal motions. The hydrodynamic parameters and predicted R2/R1 ratios were calculated with HYDRONMR ^45^ from the atomic coordinates of VHH_WT taken from the X-ray structure of its complex with Vsgi4 (5IMM).

### FoldX modelling and analysis of Vsig4 top VHHs

Sidechains in all nine available structures (PDBID: 5IMM and 5IMO for VHH_mVsig4, and 5IMK, 5IML, 9EZU, 9EZV, 9EVW, 9EZH, and 9EZI for VHH_hVsig4) were initially optimized by using the FoldX command *RepairPDB*. Structure models of all VHH candidates including VHH_WT, in complex with their respective target were generated using the FoldX *BuildModel* function, by first mutating the explored residues to Alanine and then mutating to the corresponding amino acids. The interaction energy (ΔGinteraction) and VHH individual energy (ΔGVHH) were calculated using FoldX *AnalyseComplex* function and visualized in GraphPad Prism. Triplicate modelling was performed for each complex. Differences between the candidates and VHH_WT within each group were analyzed using two-way ANOVA in GraphPad Prism.

### NGS of Vsig4 libraries

Library plasmids from the input, all rounds of selection output, and the empty control (none-target selection) outputs generated in parallel with each round of selection, were used to perform NGS sample preparation. In brief, 0.5ng of plasmid template was used in a 25 µL first-round of PCR reaction following the guidelines of KAPA HiFi HotStart ReadyMix PCR Kit (Roche, KK2602). During the first round, gene of interests (GOIs, sequence from CDR2 to CDR3) were enriched, and unique molecule identifiers (UMIs) were introduced. The second round of PCR, conducted at VIB Nucleomics Core (Leuven, Belgium) incorporated the sequencing adapters, and the samples were subsequently pooled and sequenced on AVITI 2×300 Medium (PE 250) with 5% PhiX. Raw data with around 30 millions of reads from the input, 10-20 millions of reads from R1 selection outputs and 1-2 millions of reads from R2-R4 selection outputs were acquired, demultiplexed, and initially processed with Galaxy website. Paired reads were merged using PEAR, and only reads with the correct length were retained. Additionally, UMI information were used to deduplicate. DNA sequences from CDR2 to CDR3 were extracted and translated to protein sequences. The count and enrichment factor (ratio of counts in the target selection output to counts in the empty control output; 1 was used if the count in the empty control output is 0) of CDR2 and CDR3 sequences from the input and R1 outputs were analyzed separately. Enriched CDR designs were filtered from R1 selection for both mVsig4 and hVsig4 based on specific criteria (count > 10, enrichment factor > 10). An equivalent number of negative CDR designs were identified, defined as sequences present in the input but absent or minimally represented in the R1 selection outputs. Plots of the counts and enrichment of CDR sequences as well as CDR sequence logos, were generated using R.

### FoldX modelling and analysis of NGS enriched VHH sequences

Sidechains in all nine available structures (PDBID: 5IMM and 5IMO for VHH_mVsig4, and 5IMK, 5IML, 9EZU, 9EZV, 9EVW, 9EZH, and 9EZI for VHH_hVsig4) were initially optimized by using the FoldX command *RepairPDB*. Structure models incorporating enriched or negative CDR sequences were generated using the FoldX *BuildModel* function, by first mutating the explored residues to Alanine and then mutating to the corresponding amino acids. The interaction energy change (ΔΔGinteraction) and VHH individual energy change (ΔΔGVHH) were calculated using FoldX *AnalyseComplex* function. Differences between the models with enriched sequences and models with negative sequences were analyzed using one-way ANOVA in GraphPad Prism.

## Data availability

Crystallographic coordinates and final structure factors have been deposited in the Protein Data Bank (PDB) with access codes: 9EZU (hVSIG4-VHH_h1), 9EZV (hVSIG4- VHH_h2), 9EZW (hVSIG4-VHH_h3), 9EZH (hVSIG4-VHH_h4) and 9EZI (hVSIG4-VHH_h5).

The assigned chemical shift lists were deposited in Biological Magnetic Resonance Data Bank (BMRB) entries 52361 (VHH_WT) and 52360 (VHH_h4).

## Code availability

The code written to perform the sequence design by EvolveX is available at https://github.com/SwitchLab-VIB/EvolveX/

## Author contributions

Conceptualization: RvdK, SNS, MD, JS, LSP, FR

Methodology: RvdK, ZZ, IM, JDB, DC, ANV, SNS, MD, JS, LSP, FR Investigation: RvdK, ZZ, IM, DV, TG, KM, JDB, DC, GO, CC, ANV

Visualization: RvdK, ZZ, IM, ANV

Funding acquisition: RvdK, JS, FR, LSP, SNS, MD, NG Project administration: RvdK, JS, FR, LSP, SNS, MD Supervision: RvdK, JS, FR, LSP, SNS, MD

Writing – original draft: RvdK, ZZ

Writing – review & editing: RvdK, ZZ, IM, DV, TG, KM, JDB, DC, GO, CC, ANV, SNS, MD, JS, LSP, FR

---

**Supplementary Fig.1.**
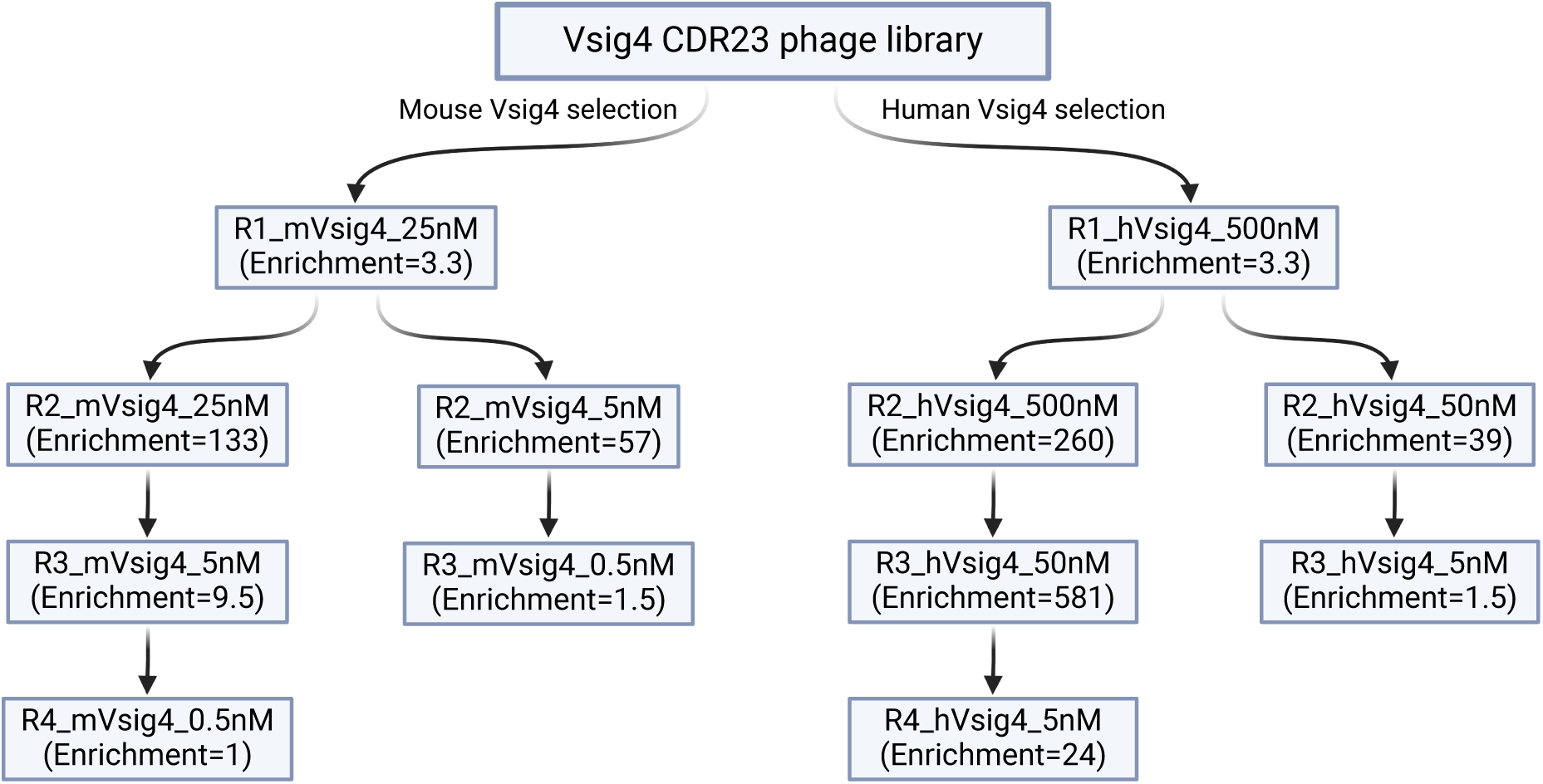
Phage library selection rounds for the combined Vsig4 CDR23 library. Graphical designs were created with BioRender.com.

**Supplementary Fig.2.**
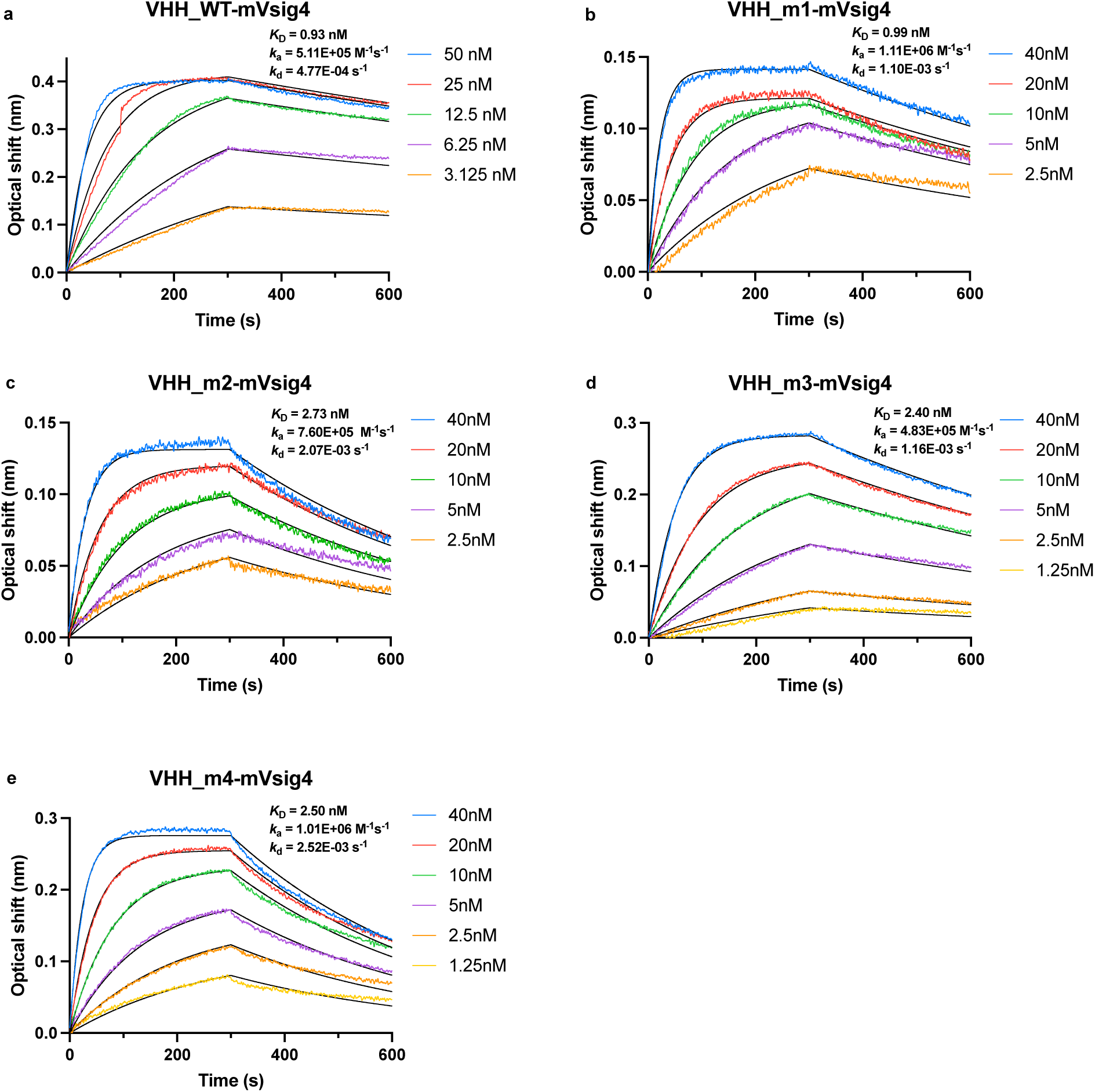
The binding curves (colored) of VHH_WT, VHH_m1-VHH_m4 against mVsig4 (BLI) with fitting curves (black).

**Supplementary Fig.3.**
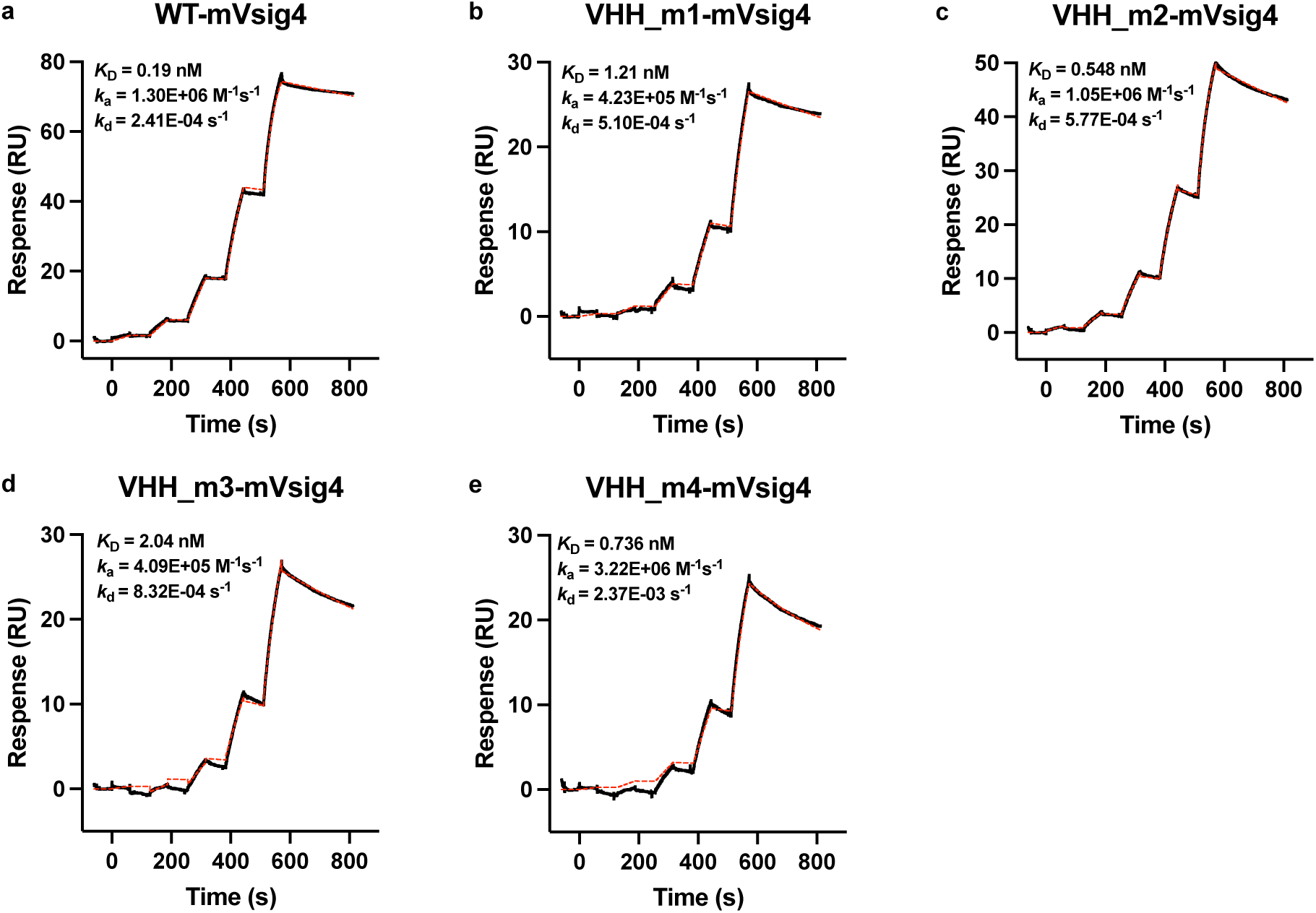
The binding curves (black) of VHH_WT, VHH_m1-VHH_m4 against mVsig4 (SPR) with fitting curves (red).

**Supplementary Fig.4.**
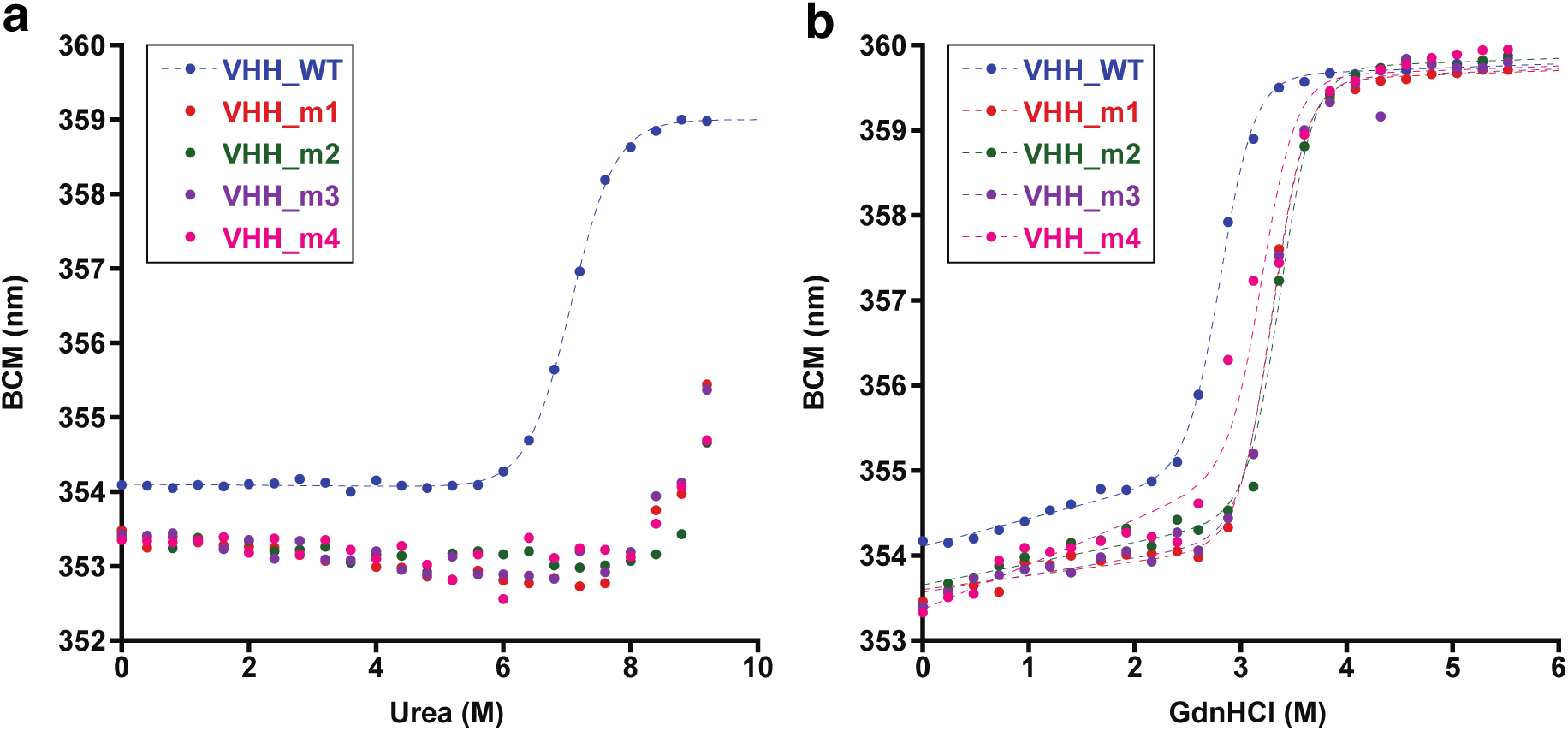
Representative chemical denaturation curves of VHH_WT, VHH_m1- m4. (**a**) Urea denaturation. (**b**) GndHCl denaturation. Fits were done using Kaleidagraph. Average ΔG is listed in Supplementary Table.3.

**Supplementary Fig.5.**
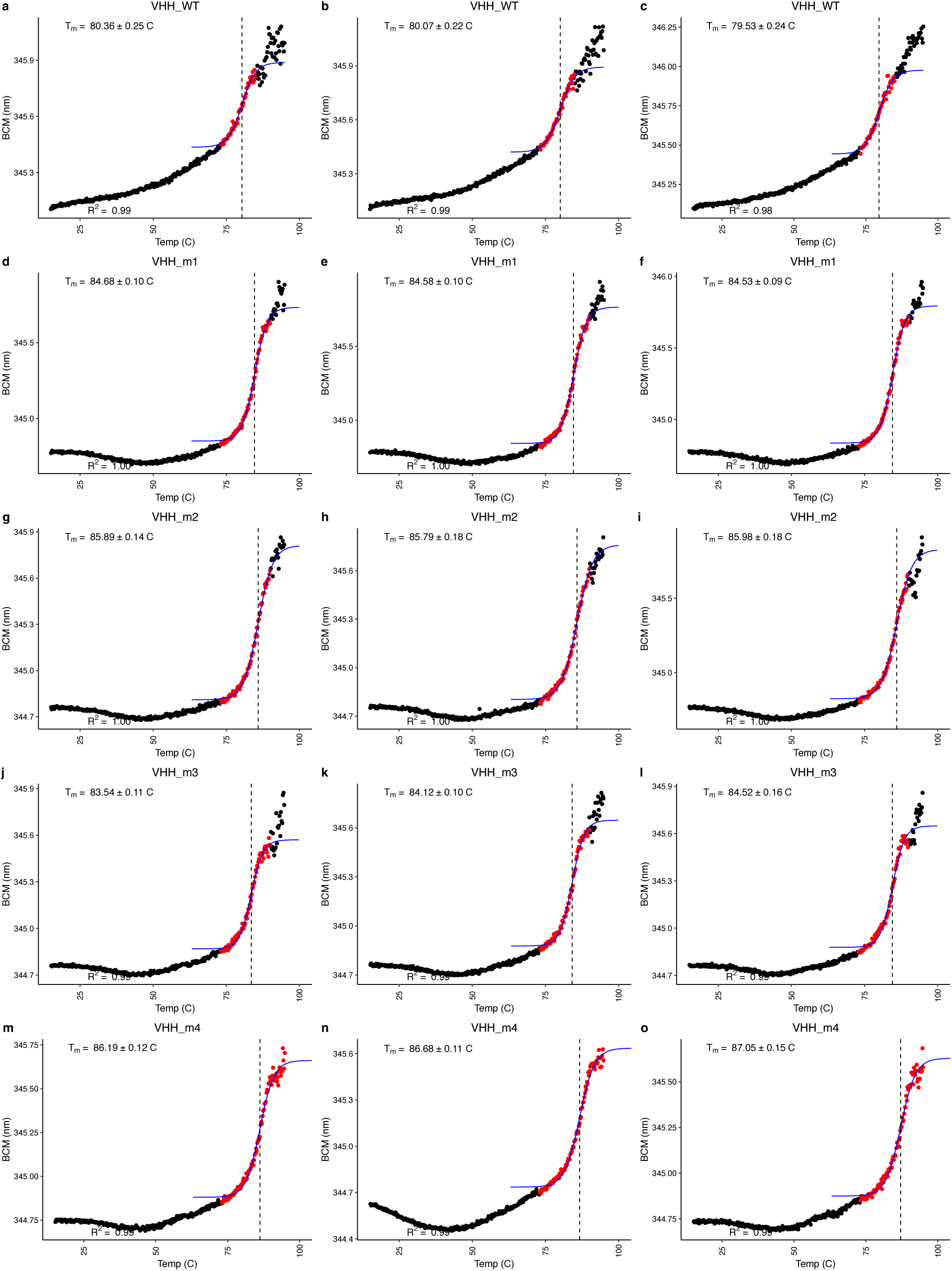
Tm determination based on Barycentric Mean calculation from Intrinsic Tryptophan Fluorescence (ITF) during a temperature ramp for VHH_WT, VHH_m1-VHH_m4.

**Supplementary Fig.6.**
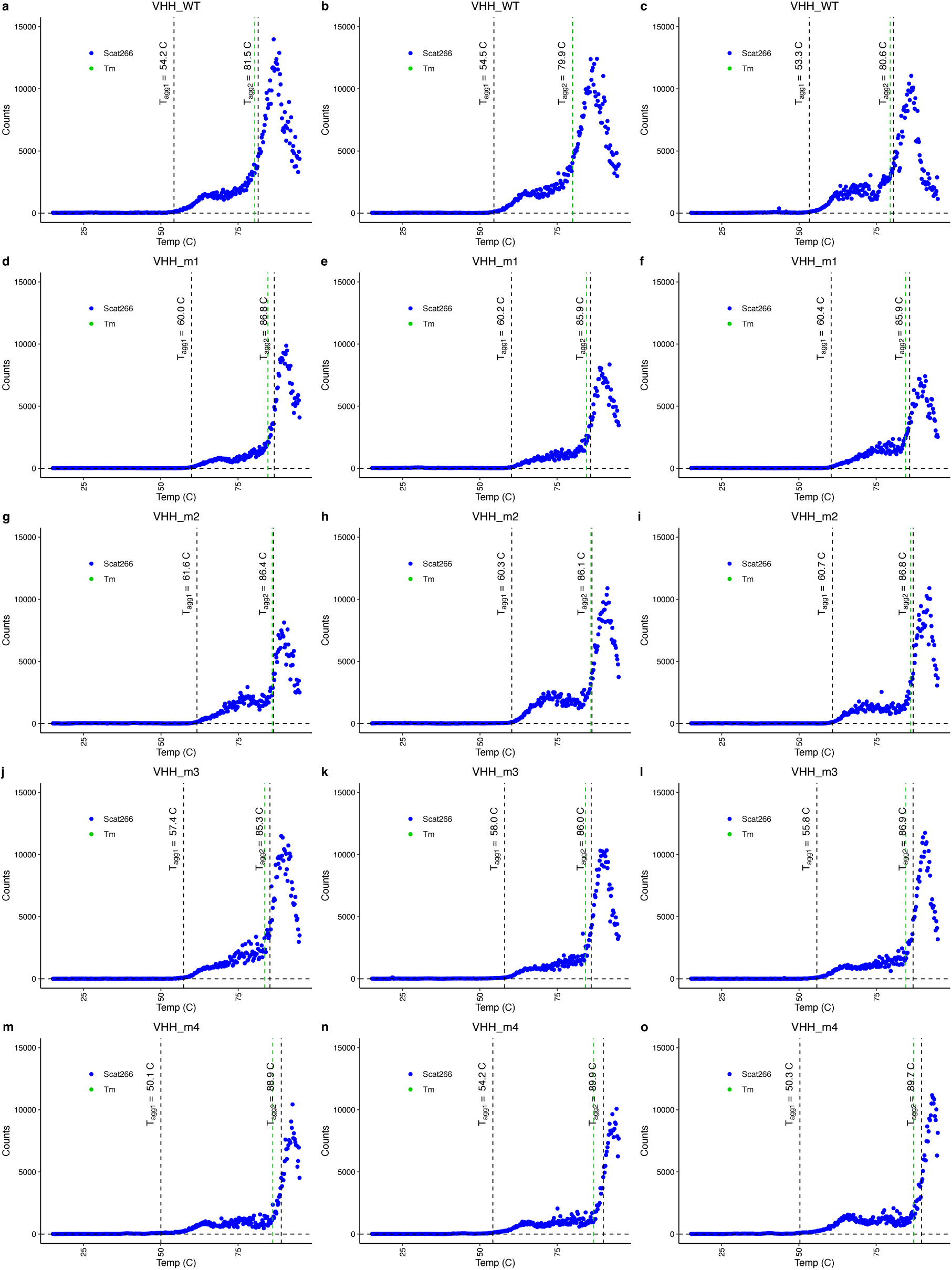
T_agg_ determination based on Scatting at 266 nm from Right Angle Light Scattering (RALS) during a temperature ramp for VHH_WT, VHH_m1-VHH_m4

**Supplementary Fig.7.**
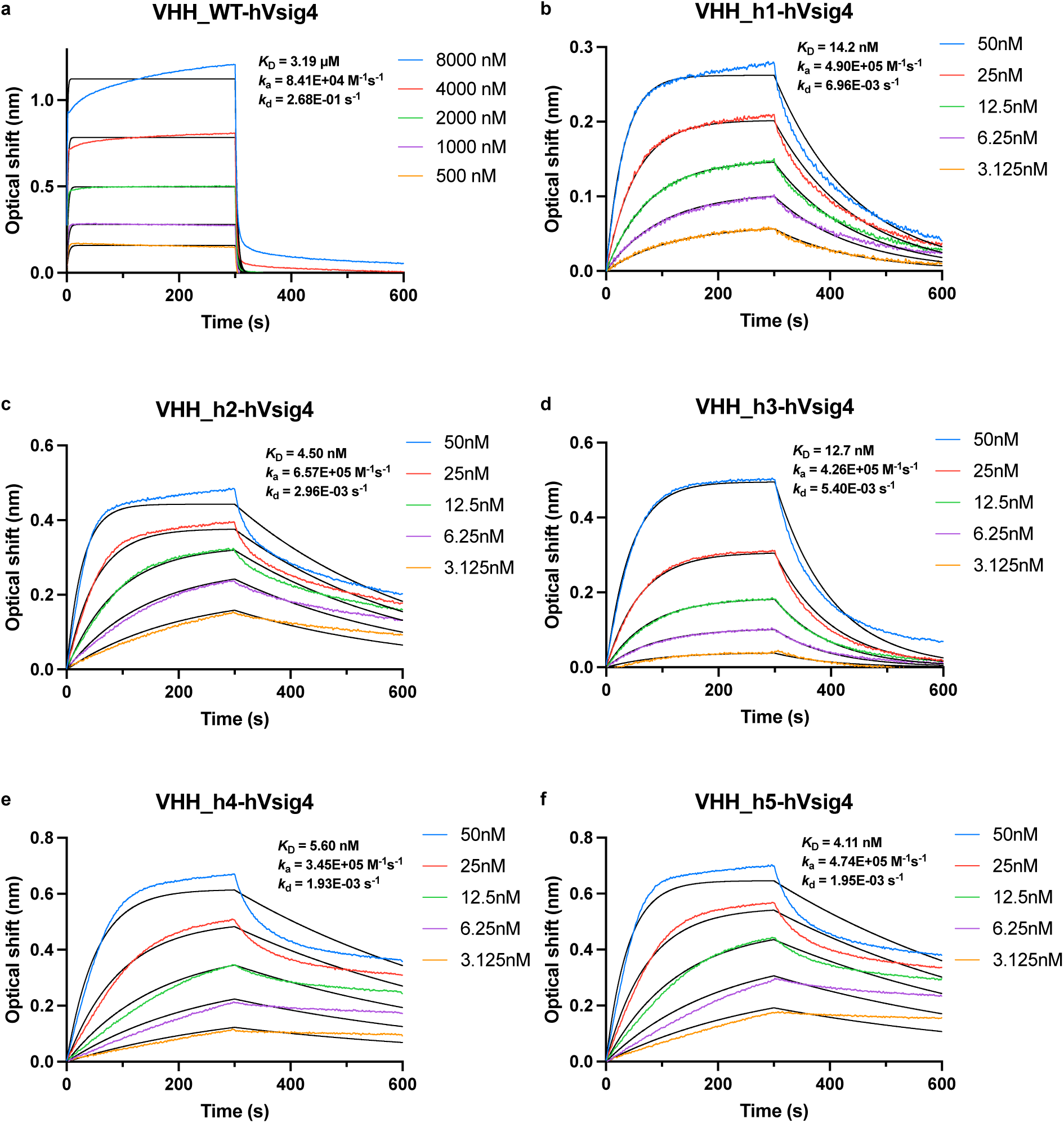
The binding curves (colored) of VHH_WT, VHH_h1-VHH_h5 against hVsig4 (BLI) with fitting curves (black).

**Supplementary Fig.8.**
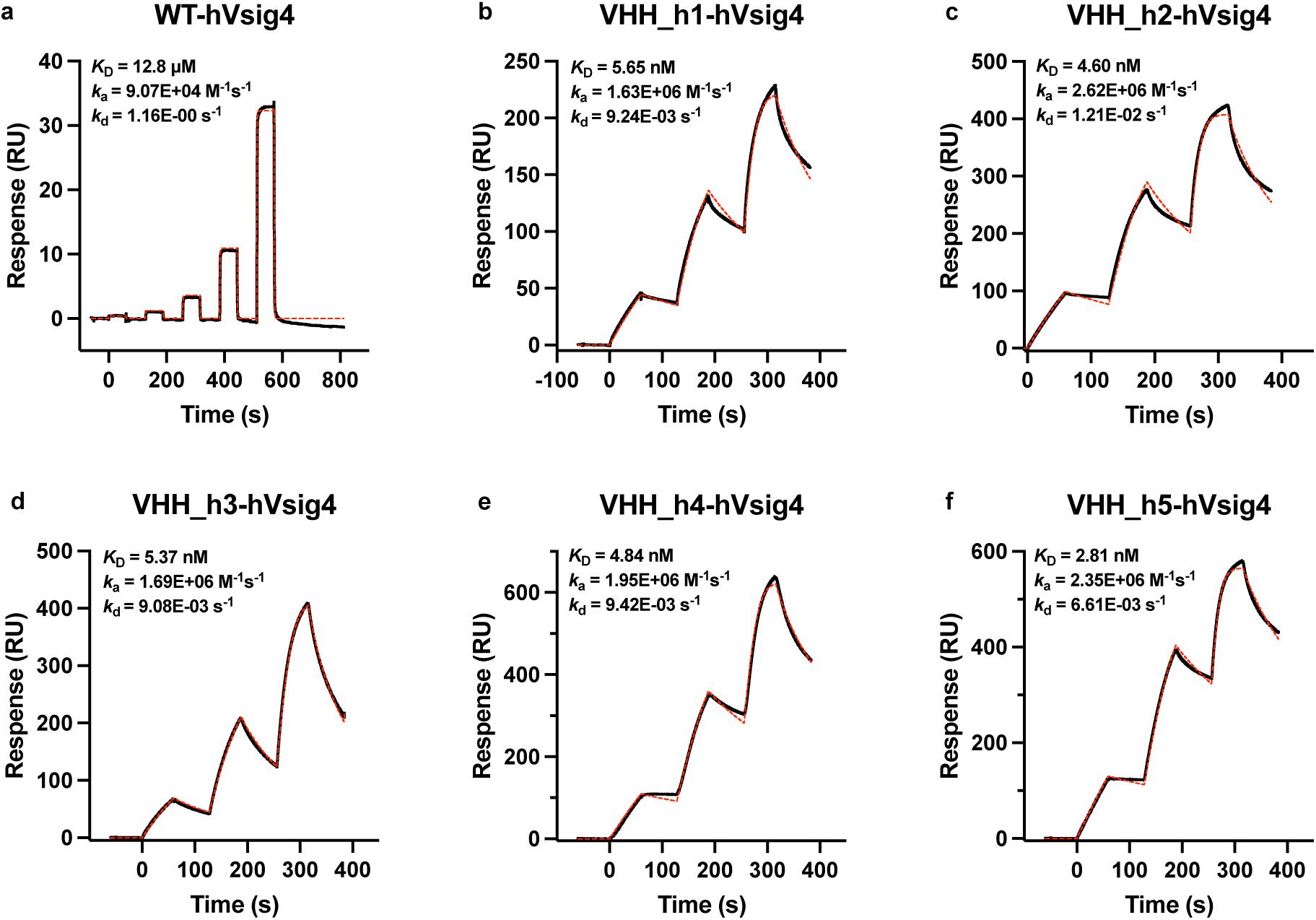
The binding curves (black) of VHH_WT, VHH_h1-VHH_h5 against hVsig4 (SPR) with fitting curves (red).

**Supplementary Fig.9.**
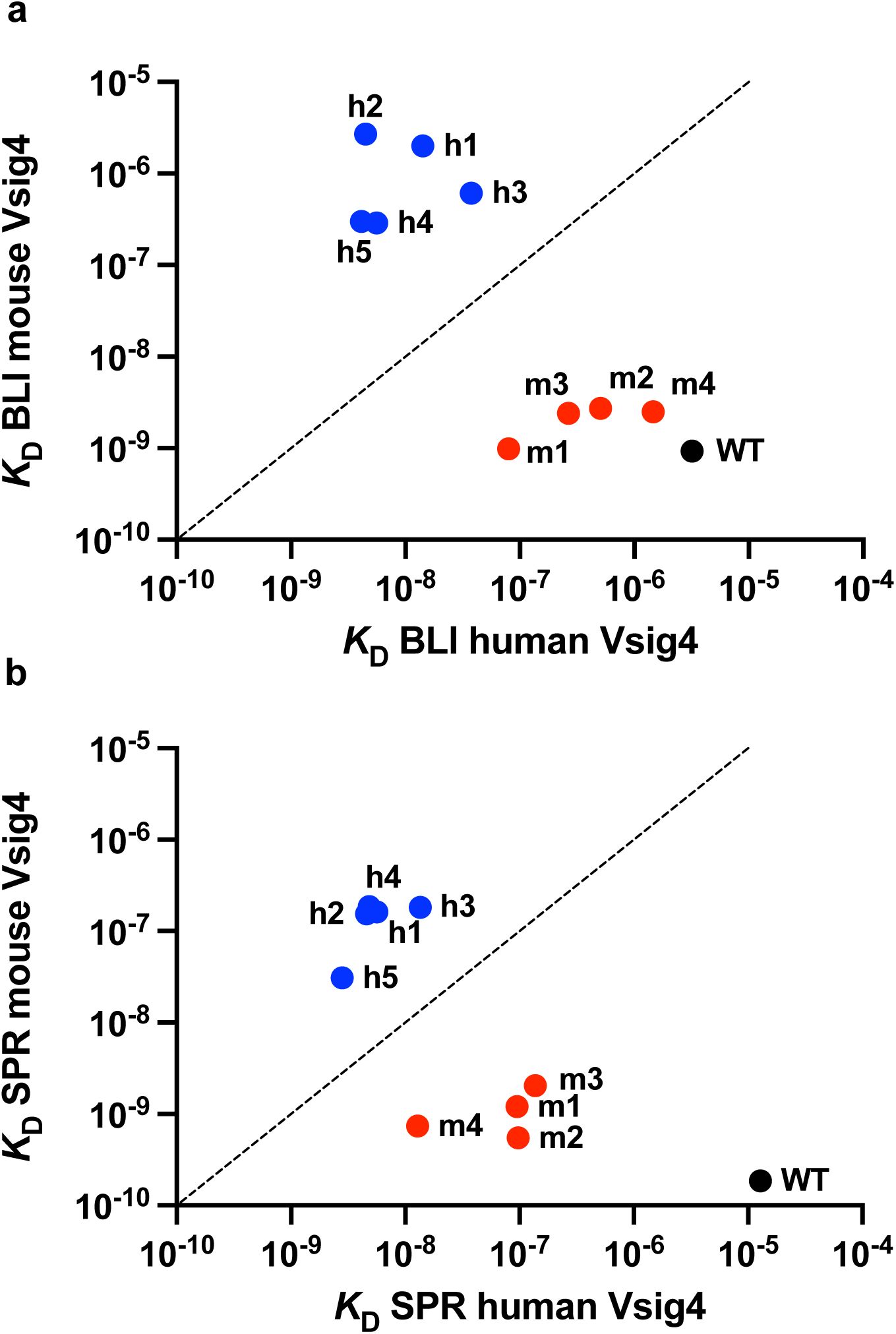
Binding of VHH_WT, VHH_m and VHH_h hits to mVsig4 and hVsig4. (**a**) Affinity constants measured by BLI of all Vsig4 hits to both mVsig4 and hVsig4. (**b**) Affinity constants measured by SPR of all Vsig4 hits to both mVsig4 and hVsig4.

**Supplementary Fig.10.**
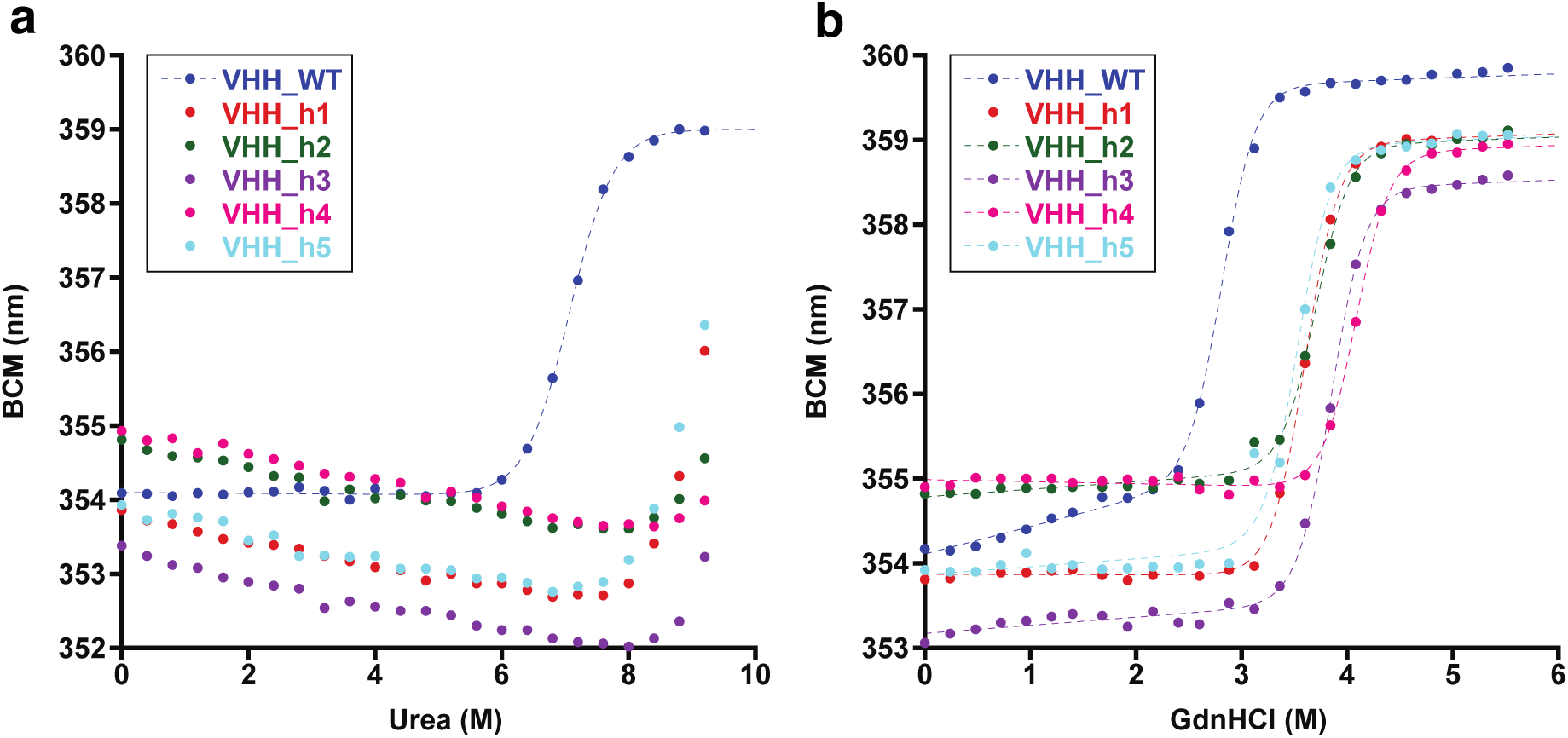
Representative chemical denaturation curves of VHH_WT, VHH_h1-h5. (**a**) Urea denaturation. (**b**) GndHCl denaturation. Fits were done using Kaleidagraph. Average ΔG is listed in **Supplementary Table.6**.

**Supplementary Fig.11.**
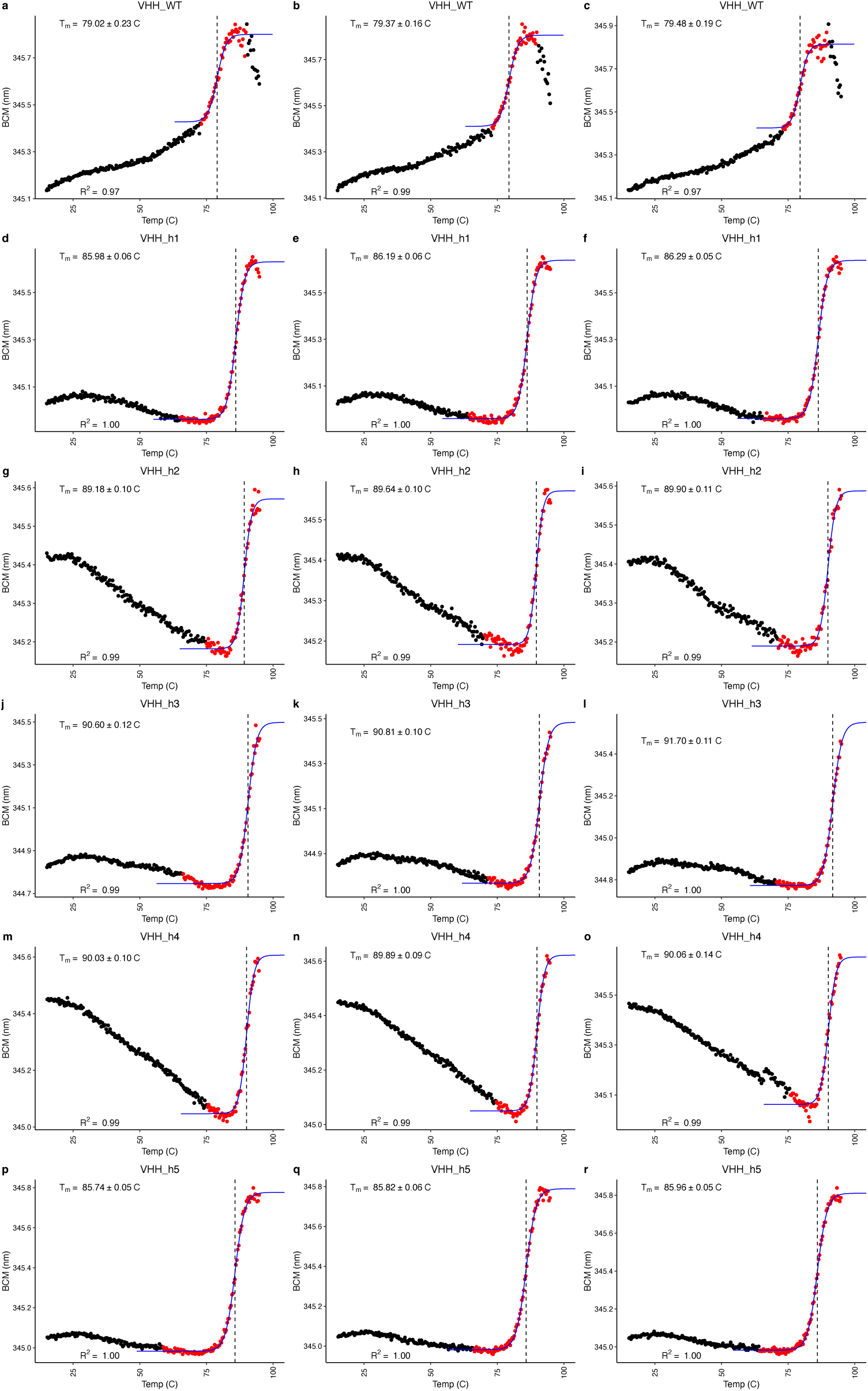
Tm determination based on Barycentric Mean calculation from Intrinsic Tryptophan Fluorescence (ITF) during a temperature ramp for VHH_WT, VHH_h1-VHH_h5.

**Supplementary Fig.12.**
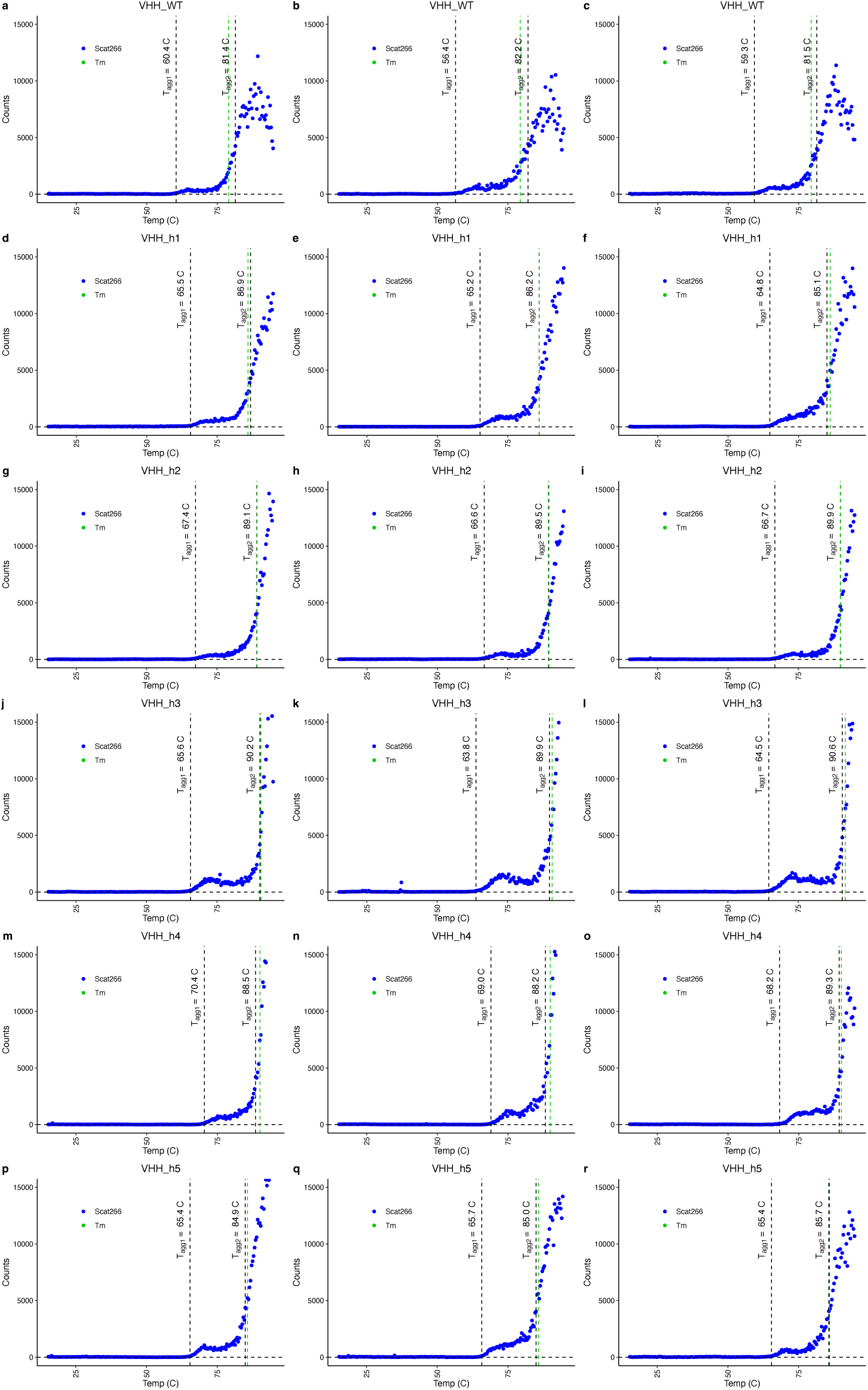
T_agg_ determination based on Scatting at 266 nm from Right Angle Light Scattering (RALS) during a temperature ramp for VHH_WT, VHH_h1-VHH_h5

**Supplementary Fig.13.**
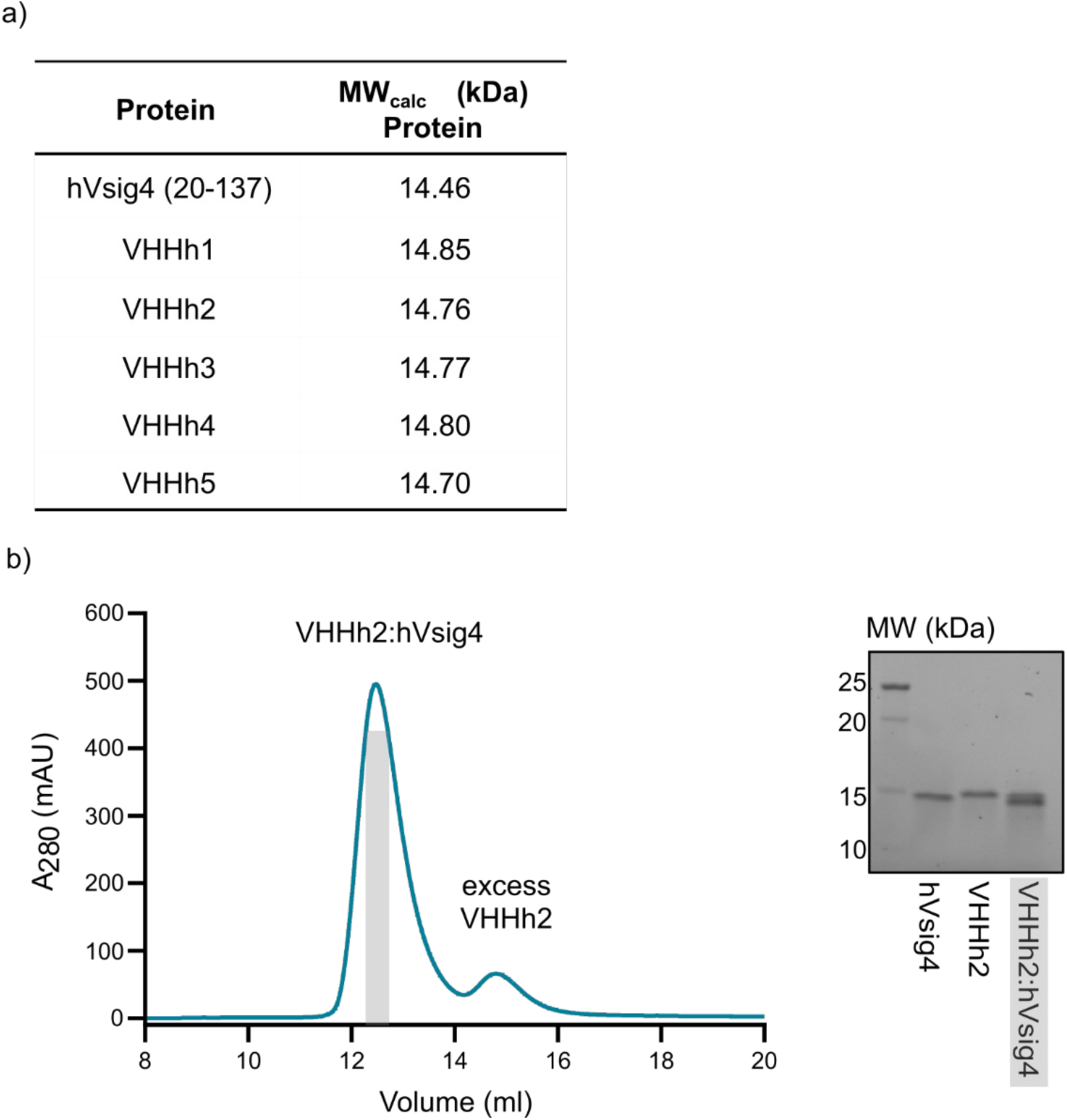
Representative SEC elution profiles for VHH_h2:hVsig4 complex reconstitution. (**a**) Overview of molecular masses of all the VHHs and hVsig4. (**b**) Representative SEC elution profile for the purification of the VHH_h2:hVsig complex using excess of VHH_h2 on a Superdex 75 Increase 10/300 GL column and representative SDS-PAGE.

**Supplementary Fig.14.**
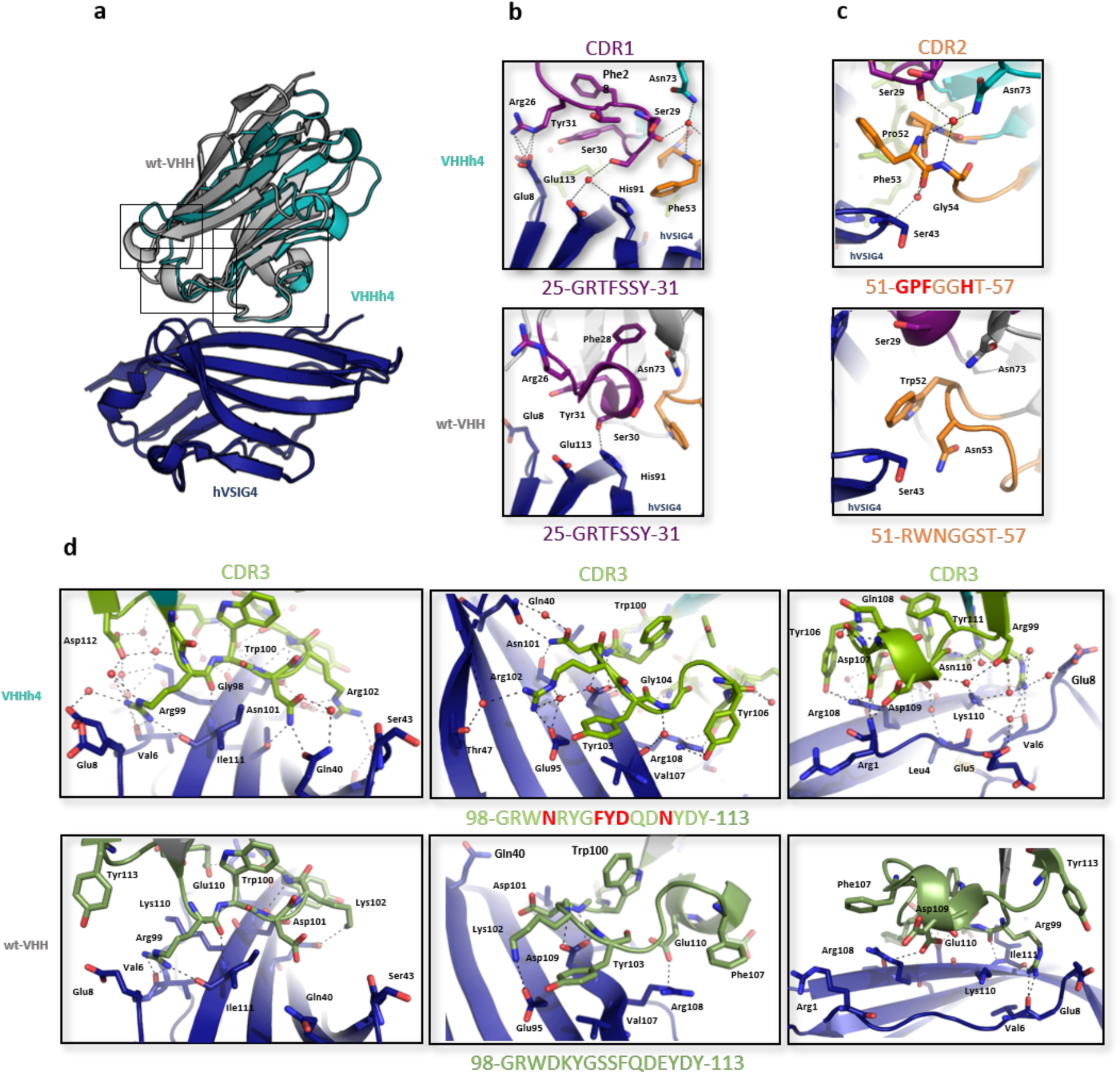
**Structure of the hVsig4:VHH_h4 complex compared to the hVsig4:VHH_WT complex**. (**a**) Cartoon representation of the determined X-ray structure for VHH_h4 fragment in complex with hVsig4 superimposed to hVsig4:VHH_WT complex (PDB 5IML) based on the structural alignment of the two hVsig4 structures. VHH_h4 is colored in teal, VHH_WT in grey and hVsig4 in dark blue. (**b**) Details of the hVsig4:VHH_h4 (up) and hVsig:VHH_WT (down) interface focusing on CDR1 loop. CDR1 loops of VHHh4 and VHH_WT are coloured in deep purple. (**c**) Details of the hVsig4:VHH_h4 (up) and hVsig:VHH_WT (down) interface focusing on CDR2 loop. CDR2 loops of VHH_h4 and VHH_WT are coloured in orange. (**d**) Details of the hVsig4:VHH_h4 (up) and hVsig:VHH_WT (down) interface focusing on CDR3 loop with 3 different perspectives. Selected interface residues are labelled and shown as sticks. CDR3 loops of VHH_h4 and VHH_WT are coloured in green (split pea). Dashed lines represent salt bridges and hydrogen bonds. Red spheres represent water molecules which are part of the interface.

**Supplementary Fig.15.**
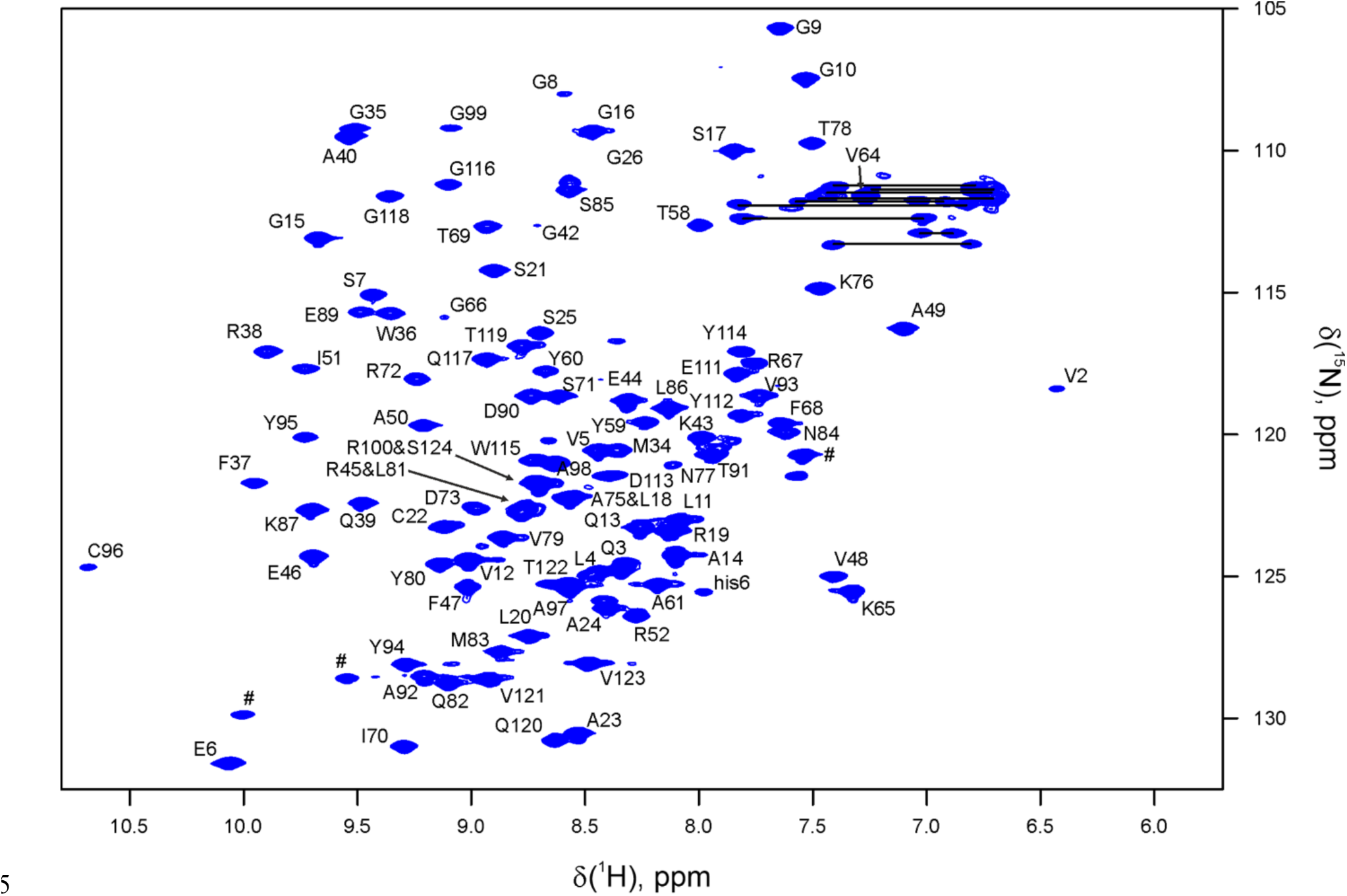
[^1^H,^15^N] HSQC spectrum of VHH_WT. The labels show assigned backbone amide resonances, tryptophan Nε1Hε1 indoles (#), a His-tag peak (his6), and sidechain NH2 resonances of Asn and Gln residues (horizontal lines). The experiments were performed in 20 mM sodium phosphate 50 mM NaCl pH 6.5 at 313 K.

**Supplementary Fig.16.**
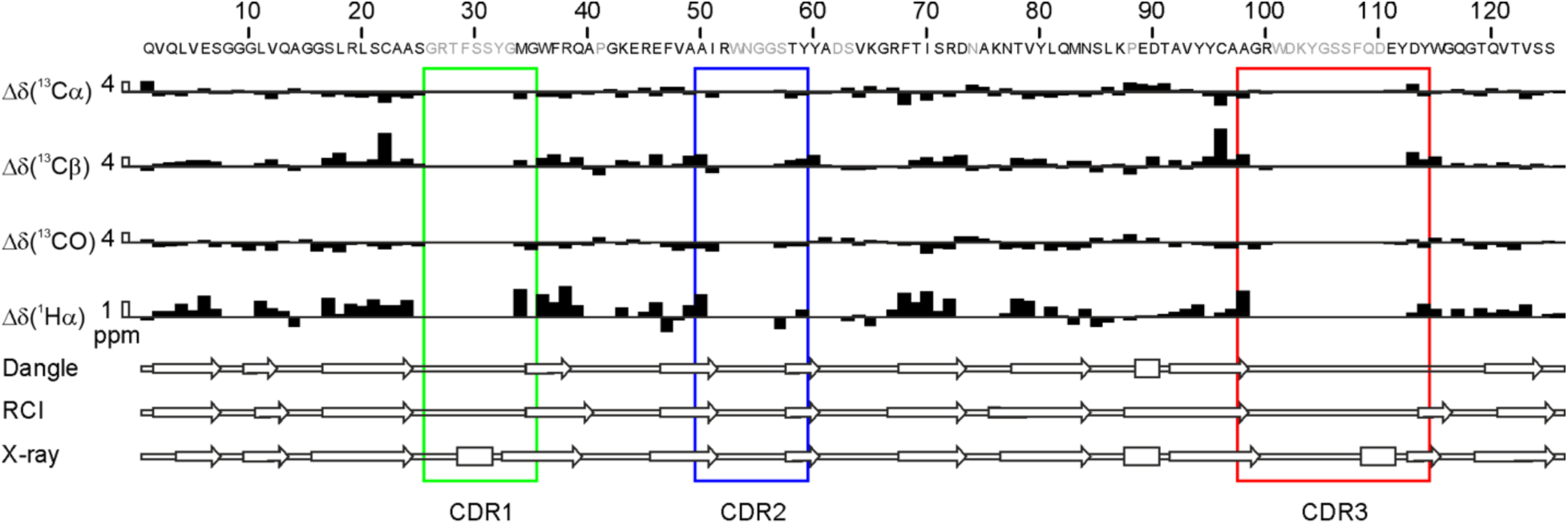
Extent of the NMR backbone assignment and secondary structure prediction of VHH_WT. Threshold deviations from random-coil ^13^Cα, ^13^Cβ, ^13^CO, and ^1^Hα chemical shifts (filled bars), with the leading open bars for scale (values in ppm). The secondary structure (rectangles – α helices, arrows – β strands) predicted by DANGLE, RCI, or seen in the X-ray structure of the VHH_WT bound to mouse Vsgi4 (5immBA). The residues that exhibited no backbone amide resonances in the HSQC spectrum are in grey.

**Supplementary Fig.17.**
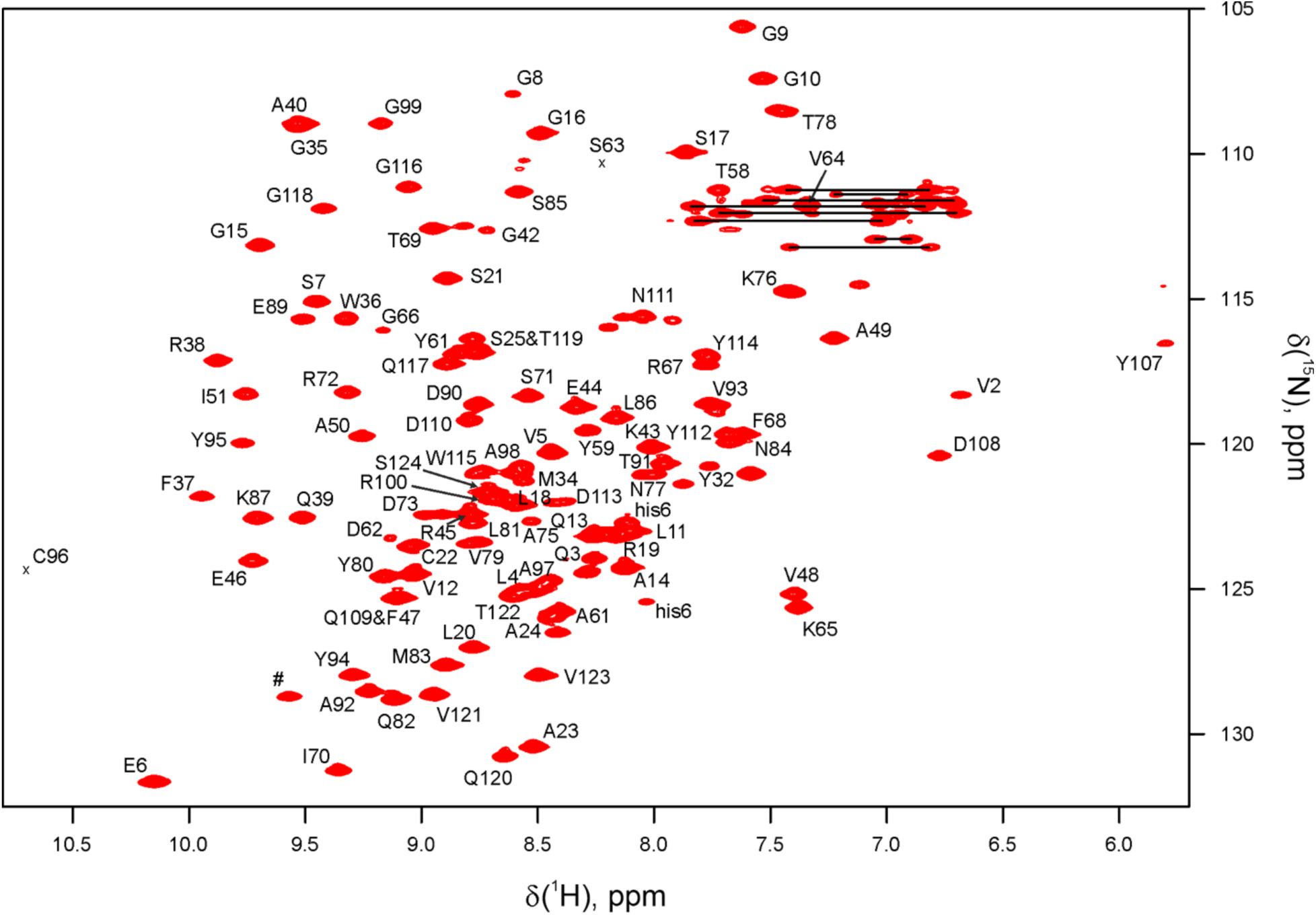
[^1^H,^15^N] HSQC spectrum of VHH_h4. The labels show assigned backbone amide resonances, tryptophan Nε1Hε1 indole (#), His-tag peaks (his6), and sidechain NH2 resonances of Asn and Gln residues (horizontal lines). Positions of S63 and C96 backbone amide signals (not seen at this contour level) are indicated by crosses. The experiments were performed in 20 mM sodium phosphate 50 mM NaCl pH 6.5 at 313 K.

**Supplementary Fig.18.**
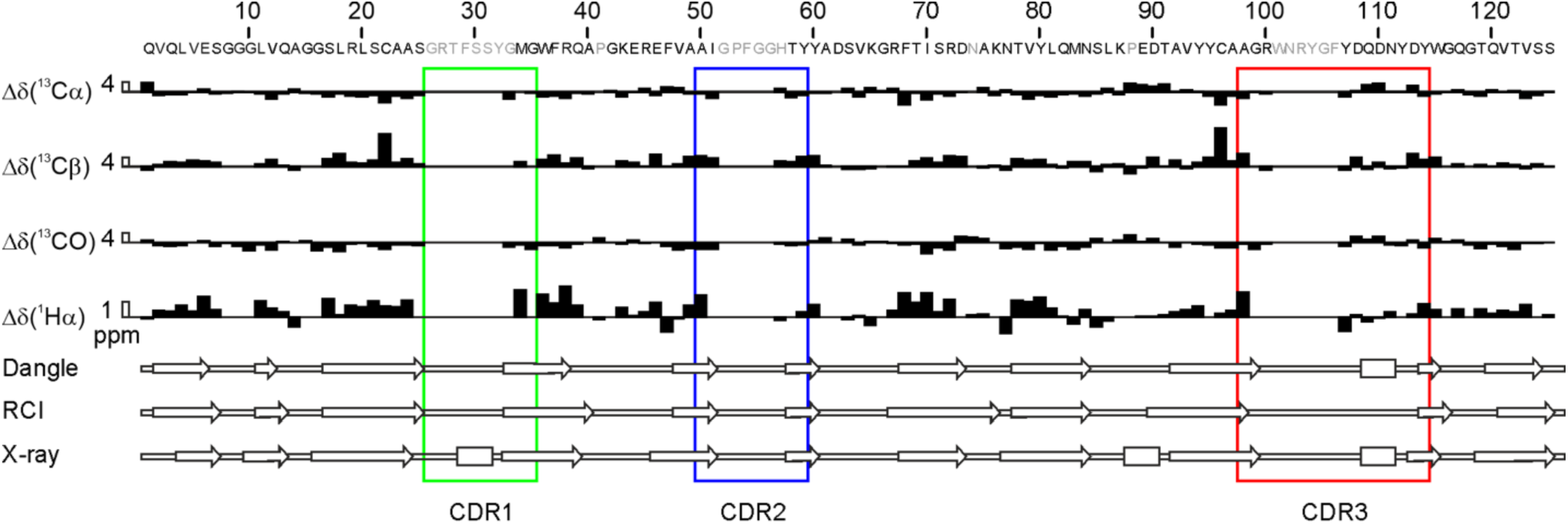
Extent of the NMR backbone assignment and secondary structure prediction of VHH_h4. Threshold deviations from random-coil ^13^Cα, ^13^Cβ, ^13^CO, and ^1^Hα chemical shifts (filled bars), with the leading open bars for scale (values in ppm). The secondary structure (rectangles – α helices, arrows – β strands) predicted by DANGLE, RCI, or seen in the X-ray structure of the VHH_WT bound to mouse Vsgi4 (5immBA). The residues that exhibited no backbone amide resonances in the HSQC spectrum are in grey.

**Supplementary Fig.19.**
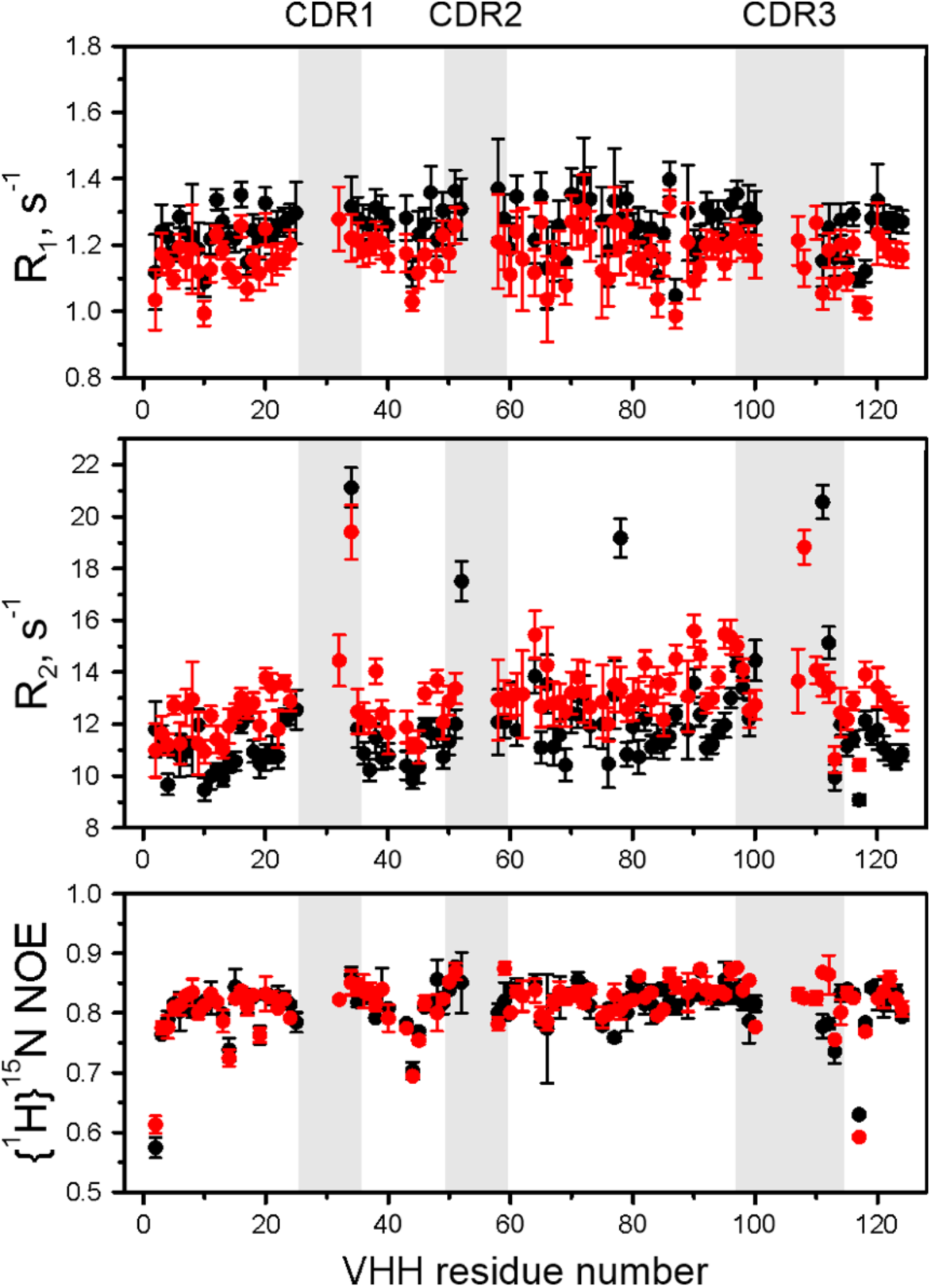
^15^N NMR relaxation parameters of VHHs. Spin-lattice (R1) and spin-spin (R2) relaxation rates, and heteronuclear steady-state nuclear Overhauser effect ({^1^H}-^15^N NOE) of the backbone amide atoms of VHH_WT (black) and VHH_h4 (red). The grey areas indicate CDR regions. Experiments were performed in 20 mM NaPi 50 mM NaCl pH 6.5 at 313 K.

**Supplementary Fig.20.**
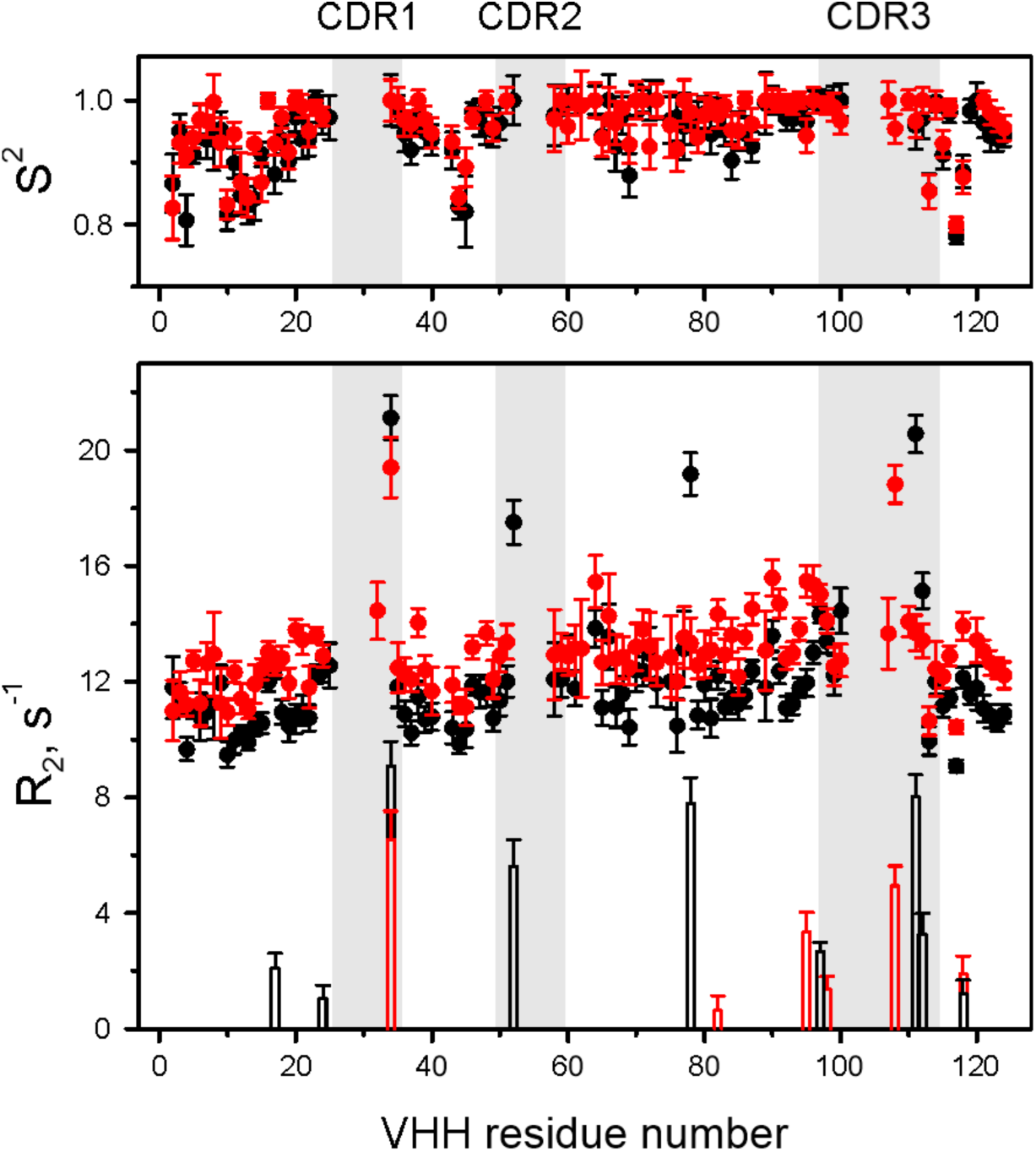
Backbone dynamics of VHHs. (top) S^2^ order parameters for the N-H vectors obtained from the model-free analysis of the NMR relaxation data in Supplementary Fig.19. (bottom) Measured R2 relaxation rates (circles) and R2,ex exchange contributions obtained from the model-free analysis (bars). The data for the VHH_WT and VHH_h4 are in black and red, respectively. The grey areas indicate CDR regions. S2 metric indicates very limited ps-ns dynamics, with the smallest values for the N-terminus and loops centered on residues 10, 44, and 117 (top). Interestingly, the few VHH_WT CDR residues that do exhibit HSQC peaks show considerable exchange contributions to transverse relaxation rates (R2,ex), suggesting that they undergo a conformational exchange on the µs-ms timescale (bottom). The R2,ex contributions are reduced in the VHH_h4 (bottom), suggesting a much less extensive conformational exchange of the mutant CDRs, which explains its more favorable spectral properties.

**Supplementary Fig.21.**
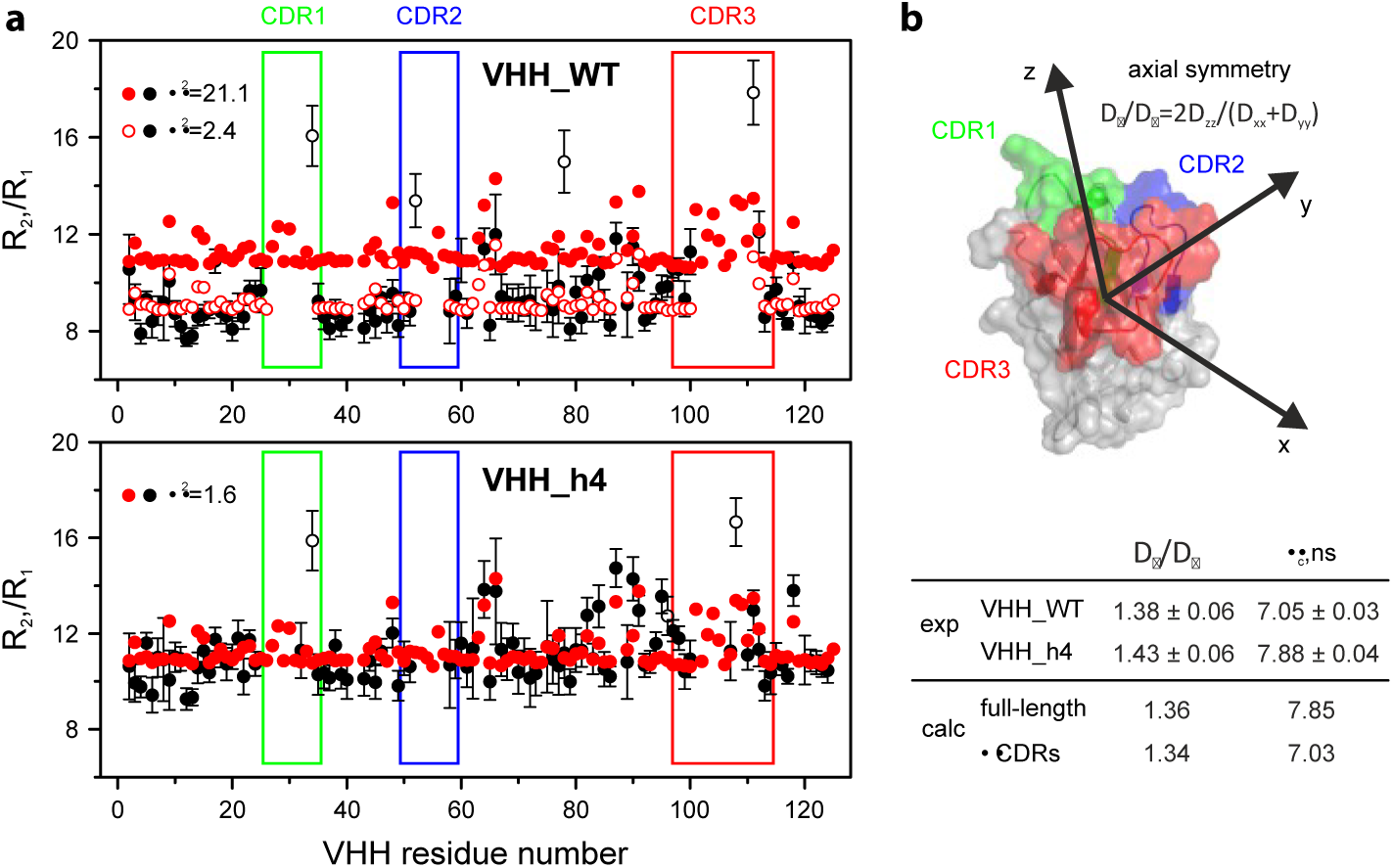
Hydrodynamic properties of VHHs. (**a**) Measured (black) and predicted (red) ratios of ^15^N R1 and R2 NMR relaxation rates for the VHH_WT (top) and VHH_h4 (bottom). R2/R1 values were calculated from the X-ray structure of the full-length VHH_WT or with the CDRs removed (filled and open red circles, respectively). VHH residues with R2 exchange contributions (black open circles) were excluded from the χ^2^ calculations. (**b**) Orientation of the diffusion tensor (coordinate system) in the VHH molecular frame. The structure of the VHH_WT is taken from the X-ray structure of its complex with the mouse Vsgi4 (5immBA), with CDRs indicated. The relaxation properties of both VHHs are well accounted for by an axially symmetric diffusion model, with the diffusion tensor parameters given in (b). Calculated from the X-ray structure of VHH bound to mouse Vsgi4, the hydrodynamic parameters of VHH_h4 are in excellent agreement with those derived from experiment: closely matching values of rotational correlation time (τc) and high similarity of experimental and calculated R2/R1 profiles (χ2 = 1.6) are particularly noteworthy (a, bottom). At the same time, the measured R2/R1 and experimentally derived τc values for the VHH_WT are considerably lower than expected (a, top), indicating a faster molecular tumbling compared to VHH_h4. To simulate a molecular system where conformationally exchanging CDR loops do not contribute to the overall protein tumbling, we removed the VHH_WT CDR residues that show no HSQC peaks from the X-ray structure and re- calculated the hydrodynamic properties. Both R2/R¬ and τc parameters of this reduced set (ΔCDRs) are in excellent agreement with the experimental VHH_WT values (a, top). Overall, it appears that free VHH_h4 in solution maintains the bound-like protein conformation, with closely packed CDR loops undergoing limited motion, while the CDRs of VHH_WT are largely disordered and involved in extensive conformational exchange.

**Supplementary Fig.22.**
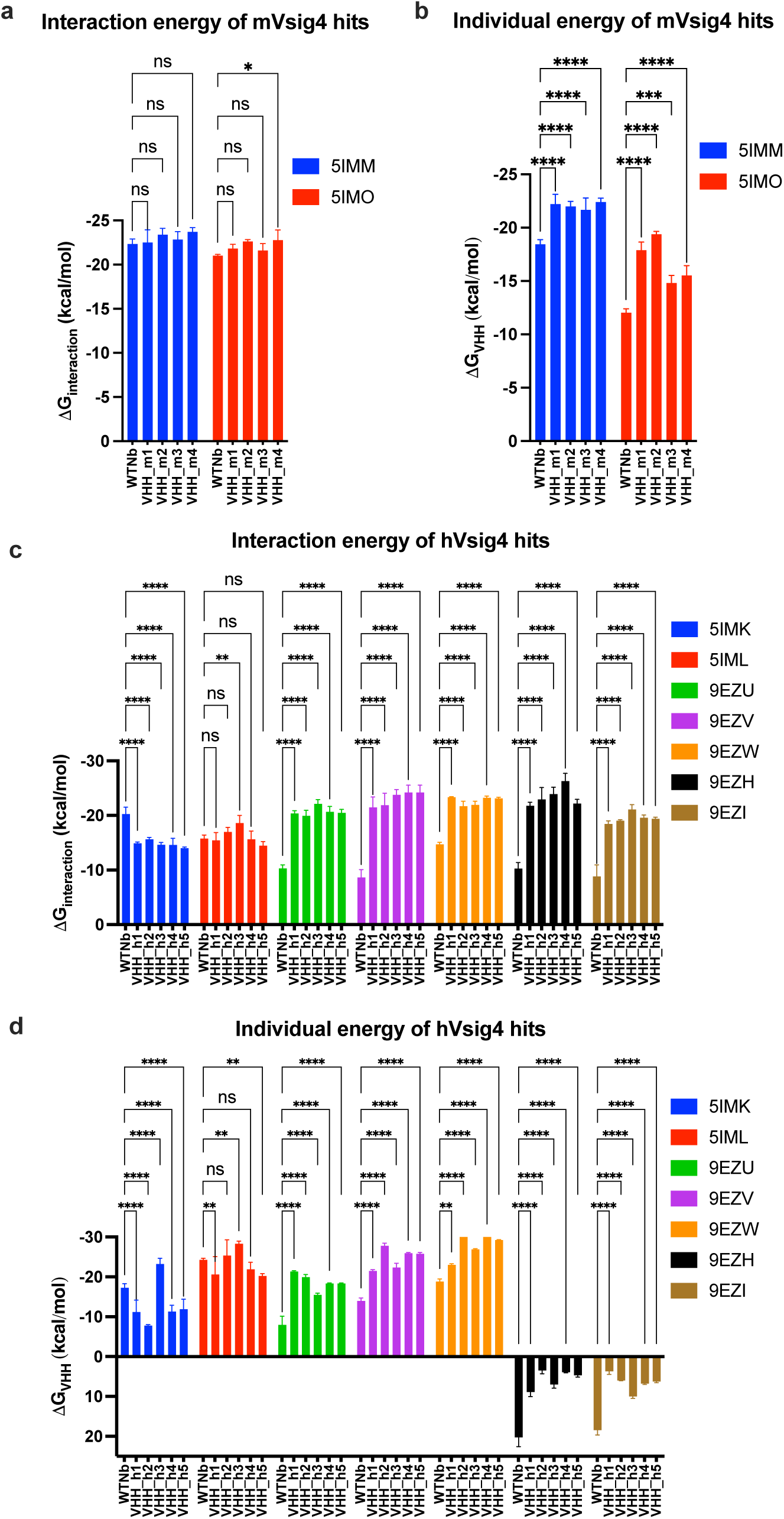
FoldX modeling of VHH hit sequences reveals energy differences between VHH_WT and VHH candidates. (**a**) Comparison of the interaction energy (ΔGinteraction) between VHH_WT and mVsig4 hits on 5IMM and 5IMO models. (**b**) Comparison of the VHH individual energy (ΔGVHH) between VHH_WT and mVsig4 hits on 5IMM and 5IMO models. (**c**) Comparison of the interaction energy (ΔGinteraction) between VHH_WT and hVsig4 hits on 5IMK, 5IML, 9EZU, 9EZV, 9EZW, 9EZH and 9EZI models. (**d**) Comparison of the VHH individual energy (ΔGVHH) between VHH_WT and hVsig4 hits on 5IMK, 5IML, 9EZU, 9EZV, 9EZW, 9EZH and 9EZI models. Triplicate modelling was performed and two-way ANOVA was run between VHH_WT and VHH candidates in all four plots (ns p>0.123, *p<0.033, **p<0.002, ***p<0.0002, ****p<0.00001).

**Supplementary Fig.23.**
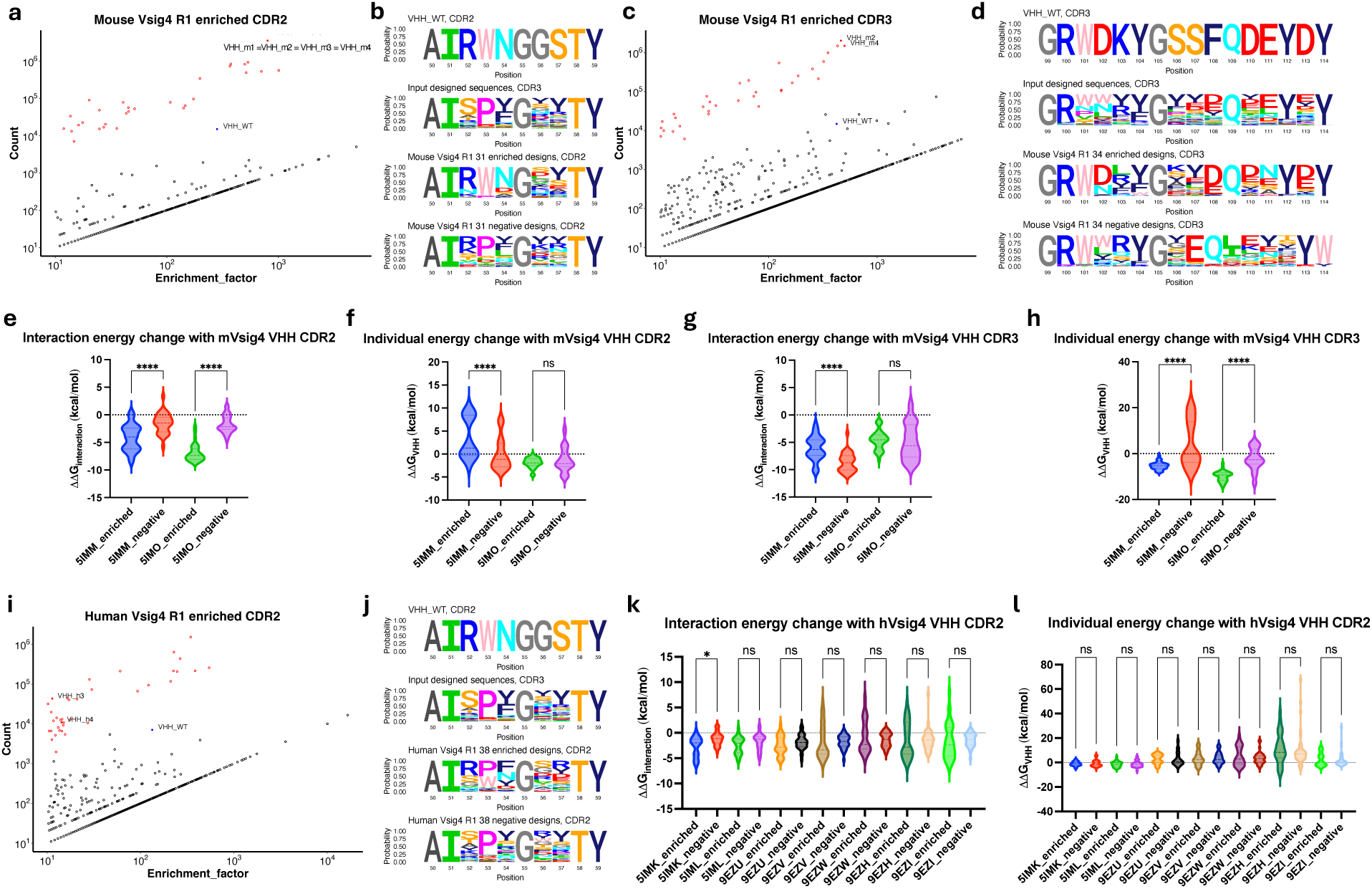
FoldX modeling of NGS-identified CDR designs reveals energy change differences between enriched and negative CDR designs. (**a**) Count and enrichment factor of mVsig4 enriched CDR2 sequences (count > 10, enrichment factor > 10, enrichment factor is the ratio of the count in target selection to the count in empty control). Red dots are the sequences designed by EvolveX, red solid dots are the sequences from identified hits, blue solid dot is VHH_WT, and black dots are non-designs or mutants generated during the PCR or phage selection stages. The same annotation is omitted in the (c) and (i) below. (**b**) Sequence logo plots of VHH_WT CDR2, 31 mVsig4 enriched CDR2 designed sequences and 31 negative CDR2 designed sequences. AbM CDR definition was used. The same annotation is omitted in the (d) and (j) below. (**c**) Count and enrichment factor of mVsig4 enriched CDR3 sequences. (**d**) Sequence logo plots of VHH_WT CDR3, 34 mVsig4 enriched CDR3 designed sequences and 34 negative CDR3 designed sequences. (**e**) Comparison of the interaction energy change (ΔΔGinteraction) between enriched and negative mVsig4 CDR2 designs. (**f**) Comparison of the VHH individual energy change (ΔΔGVHH) between enriched and negative mVsig4 CDR2 designs. (**g**) Comparison of the interaction energy change (ΔΔGinteraction) between enriched and negative mVsig4 CDR3 designs. (**h**) Comparison of the VHH individual energy change (ΔΔGVHH) between enriched and negative mVsig4 CDR3 designs. (**I**) Count and enrichment factor of hVsig4 enriched CDR2 sequences. (**j**) Sequence logo plots of VHH_WT CDR2, 38 hVsig4 enriched CDR2 designed sequences and 38 negative CDR2 designed sequences. (**k**) Comparison of the interaction energy change (ΔΔGinteraction) between enriched and negative hVsig4 CDR2 designs. (**l**) Comparison of the VHH individual energy change (ΔΔGVHH) between enriched and negative hVsig4 CDR2 designs. One way ANOVA was used in (e-h and k- l) (ns p>0.123, *p<0.033, **p<0.002, ***p<0.0002, ****p<0.00001).

**Supplementary Table.1.**
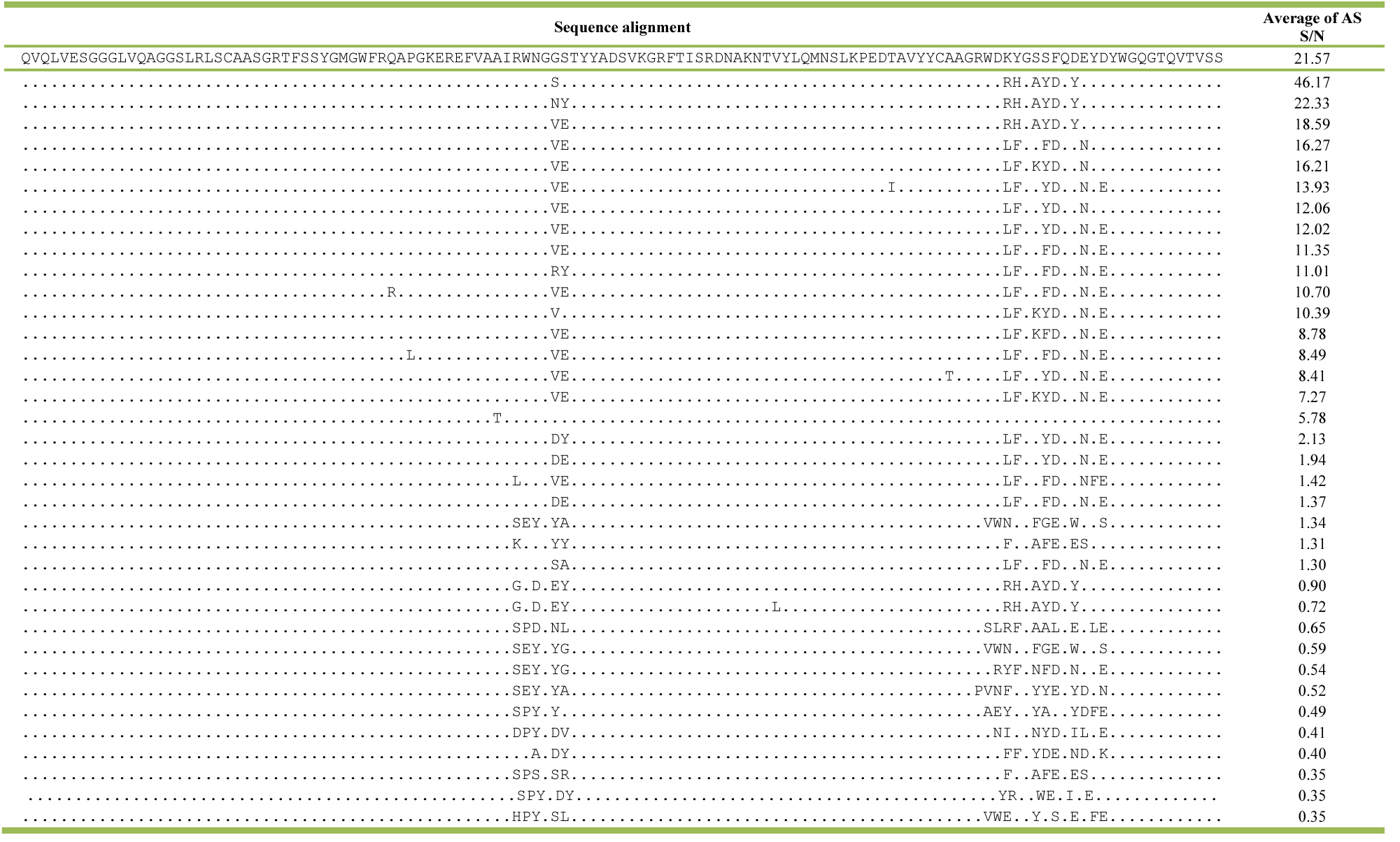
36 unique sequences of mVsig4 hits with average of AlphaScreen signal/noise (S/N) ratio.

**Supplementary Table.2.**
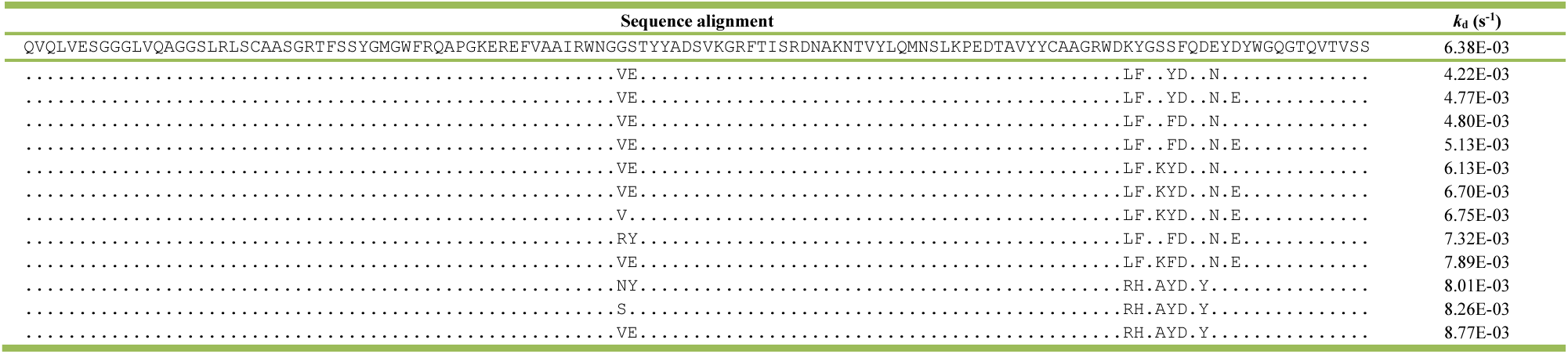
12 unique sequences of mVsig4 hits selected for the BLI off-rate test.

**Supplementary Table.3.**
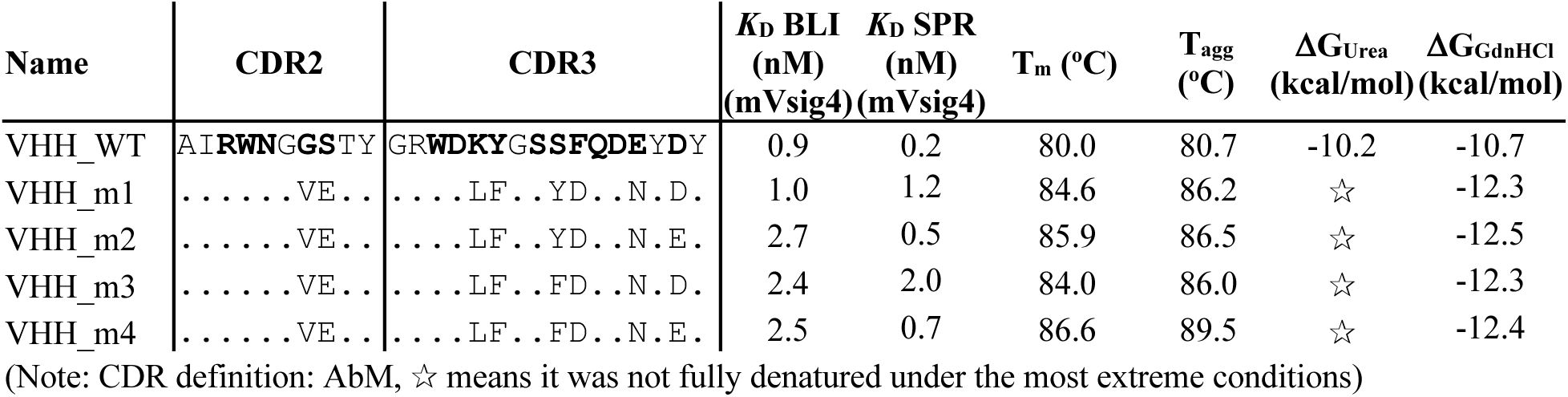
Biophysical characterization data of VHH_m1-VHH_m4.

**Supplementary Table.4.**
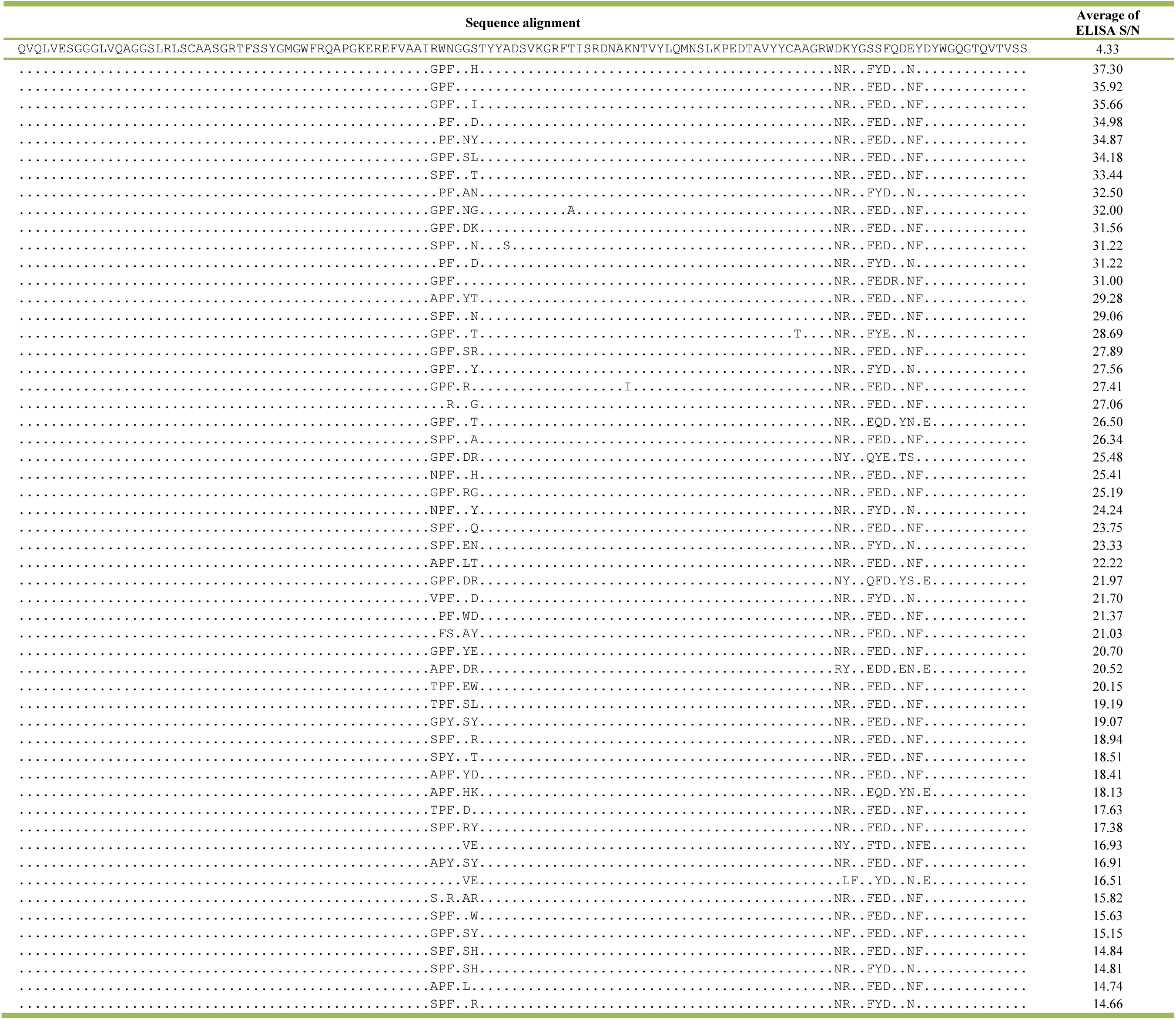
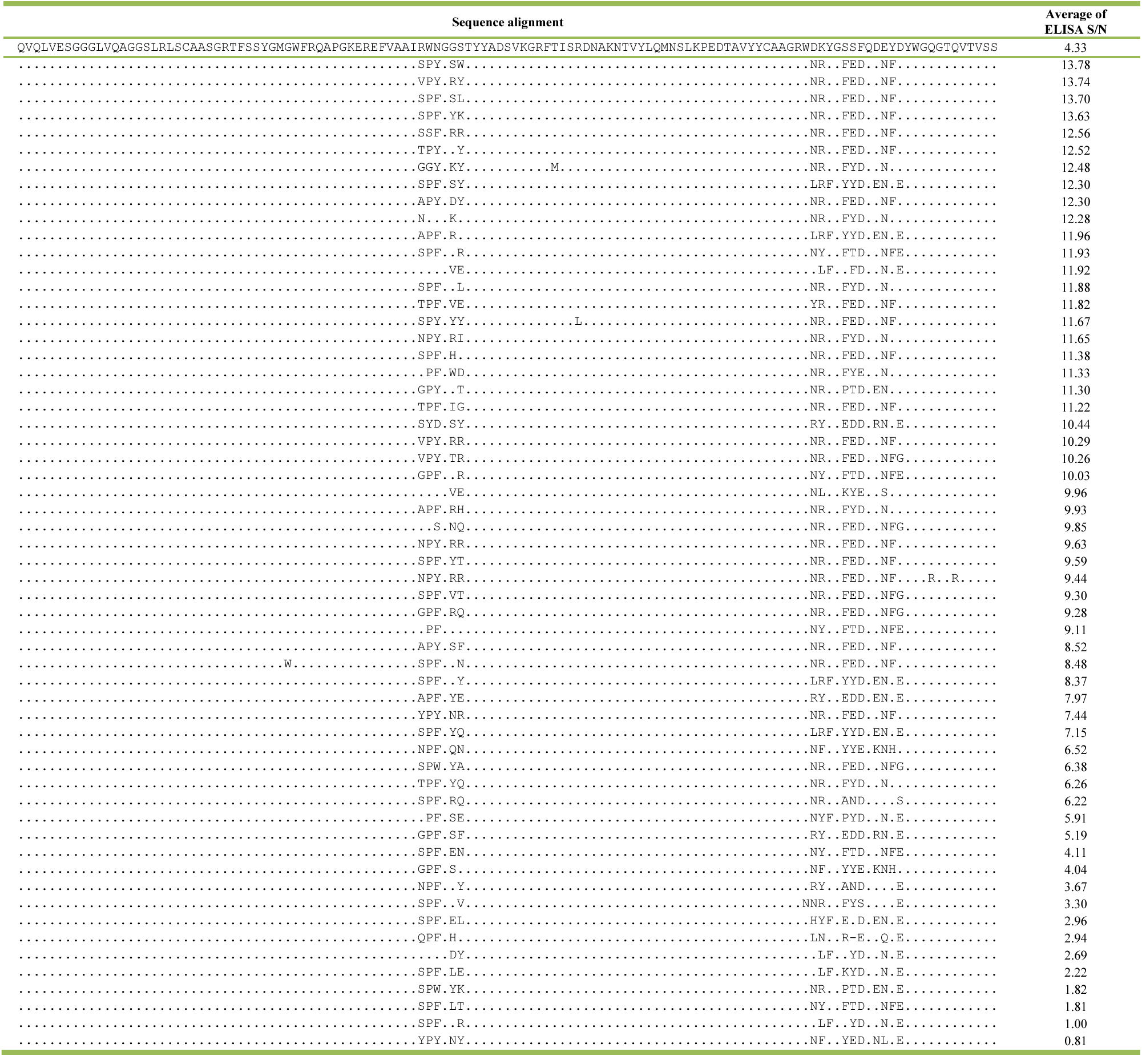
112 unique sequences of hVsig4 hits with average of ELISA signal/noise (S/N) ratio.

**Supplementary Table.5.**
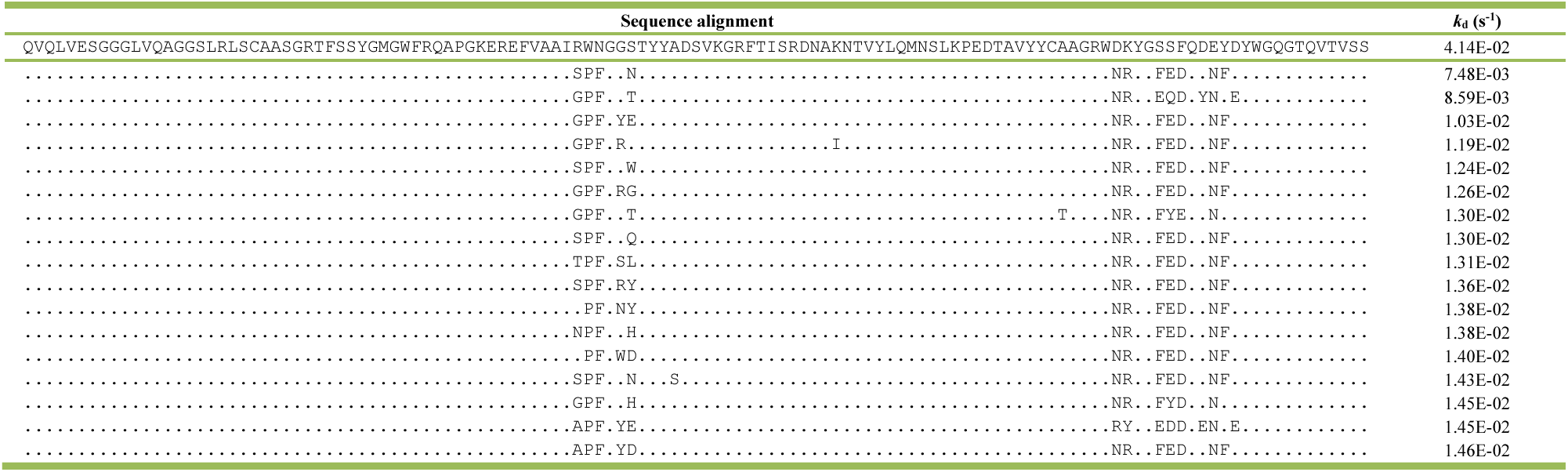
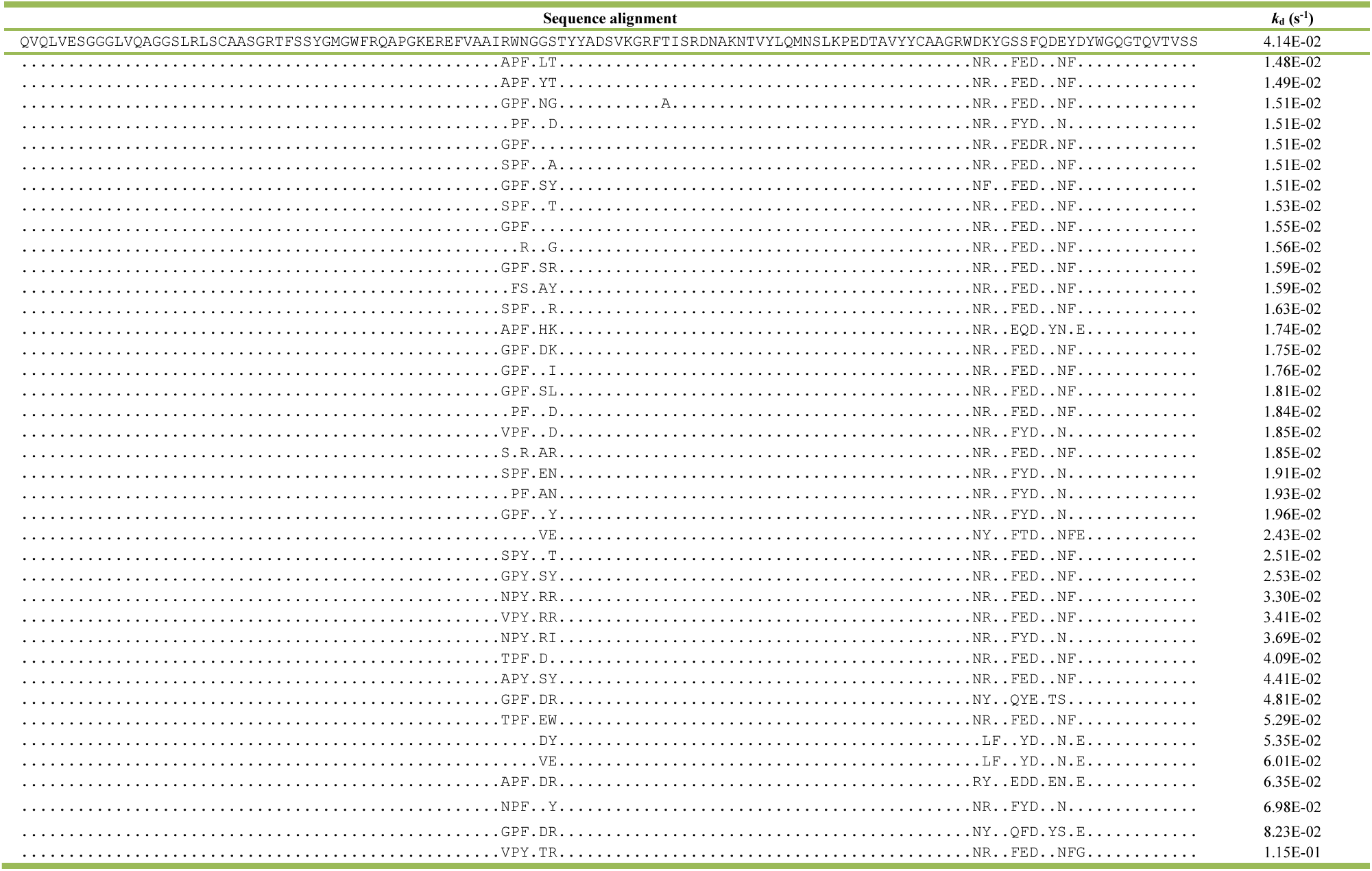
56 unique sequences of hVsig4 hits selected for the BLI off-rate test.

**Supplementary Table.6.**
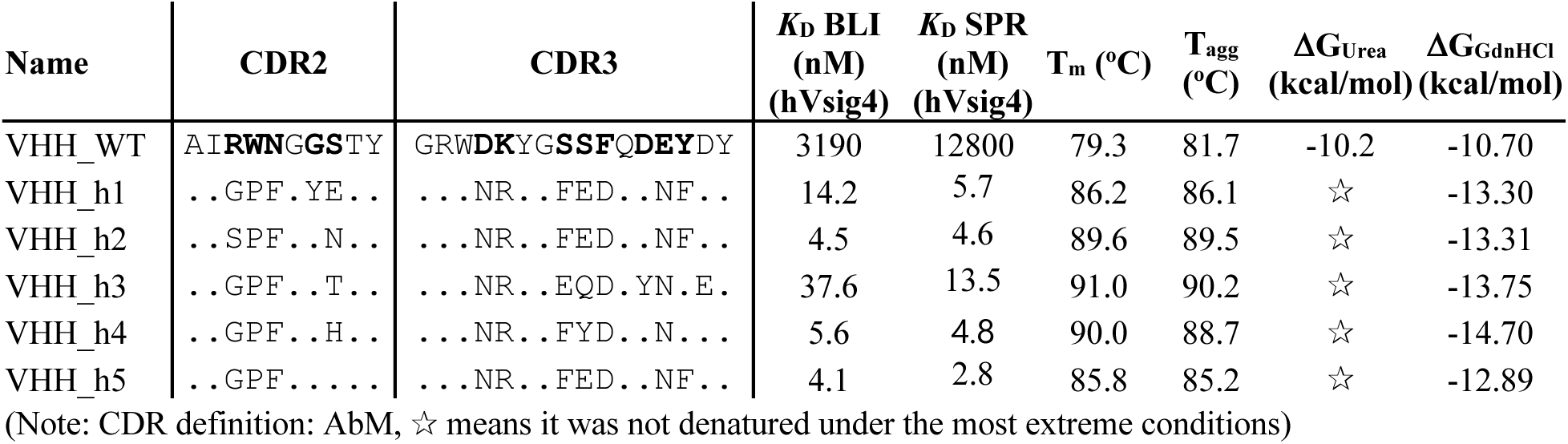
Biophysical characterization data of VHH_h1-VHH_h5.

**Supplementary Table.7.** Crystallization data for VHH_h1-5 in complex with hVsig4.

| Protein complex | hVsig4-VHH_h1 | hVsig4-VHH_h2 | hVsig4-VHH_h3 | hVsig4-VHH_h4 | hVsig4-VHH_h5 |
| --- | --- | --- | --- | --- | --- |
| Crystallization conditions | 0,17M ammonium sulfate<br>15% glycerol<br>25,5% PEG 4000 | 0,1M Na-citrate<br>pH 5,6<br>20% 2-propanol<br>21% PEG 4000 | 0,1M Na-citrate<br>pH 5,6<br>20% 2-propanol<br>21% PEG 4000 | 0,1M Bis-Tris<br>pH 6.0<br>24% PEG3350 | 0,1M Bis-Tris<br>pH 6.5<br>30% PEG3350 |
| Temperature (K) | 287 | 287 | 294 | 293 | 293 |
| Protein concentration (mg/ml) | 4,2 | 4,2 | 5,6 | 3,80 | 4,5 |
| Cryoprotectant | none | 25% ethylene glycol | 20% ethylene glycol | 15% glycerol | 15% glycerol |
| **Data collection<sup>a</sup>** |  |  |  |  |  |
| X-ray source | PETRA III (P13) | PETRA III (P13) | PETRA III (P13) | ALBA (Xaloc) | ALBA (Xaloc) |
| Space group | P2<sub>1</sub>2<sub>1</sub>2<sub>1</sub> | P2<sub>1</sub>2<sub>1</sub>2<sub>1</sub> | P2<sub>1</sub>2<sub>1</sub>2<sub>1</sub> | P2<sub>1</sub>2<sub>1</sub>2<sub>1</sub> | P2<sub>1</sub>2<sub>1</sub>2<sub>1</sub> |
| Cell dimensions |  |  |  |  |  |
| a, b, c (Å) | 50.651, 51.404, 89.649 | 52.291 53.069<br>87.531 | 60.867 129.940<br>144.280 | 25.7, 51.0, 175.2 | 25.7, 50.9, 175.9 |
| α, β, γ (°) | 90.0, 90.0, 90.0 | 90.0, 90.0, 90.0 | 90.0, 90.0, 90.0 | 90.0, 90.0, 90.0 | 90.0, 90.0, 90.0 |
| Unique reflections | 109581 (17482) | 43885 (7006) | 89487 (14083) | 18076 (928) | 17284 (825) |
| Wavelength (Å) | 0.9762 | 0.9762 | 0.9762 | 0.9793 | 0.9793 |
| Resolution (Å) | 51.40-1.05<br>(1.11-1.05) | 53.07-1.45<br>(1.54-1.45) | 129.94-1.90<br>(2.01-1.90) | 87.62-1.94<br>(1.97-1.94) | 87.95-1.97<br>(2.00-1.97) |
| R<sub>meas</sub> (%) | 10.0 (102.8) | 8.8 (218.6) | 19.8 (247.1) | 9.6 (109.4) | 11.3 (99.2) |
| <I/σ> | 12.40 (1.59) | 17.77 (1.13) | 13.61 (1.19) | 16.0 (2.5) | 13.5 (2.4) |
| Completeness (%) | 99.90 (99.42) | 99.90 (100) | 98.3 (97.1) | 100 (100) | 100 (100) |
| Redundancy | 13.3 (11.9) | 13.3 (13.3) | 13.9 (13.5) | 12.6 (12.8) | 12.5 (12.3) |
| CC (½) (%) | 99.7 (75.7) | 100 (42.3) | 99.8 (45.6) | 99.8 (91.0) | 99.8 (90.9) |
| Wilson B factor (Å<sup>2</sup>) | 15.02 | 27.10 | 36.57 | 31.67 | 31.15 |
| **Refinement<sup>b</sup>** |  |  |  |  |  |
| Resolution (Å) | 24.71 - 1.05<br>(1.06 - 1.05) | 33.56 - 1.45<br>(1.502 - 1.45) | 64.97 - 1.90<br>(1.92 - 1.90) | 44.05-1.94<br>(2.009-1.94) | 44.04-1.97<br>(2.04-1.97) |
| No. reflections | 109570 (10760) | 43876 (4327) | 89397 (8824) | 17989 (1772) | 17159 (1630) |
| R<sub>work</sub> (%) | 15.67 | 17.34 | 19.08 | 19.63 | 19.75 |
| R<sub>free</sub> (%) | 17.57 | 21.51 | 22.83 | 24.01 | 24.30 |
| No. non-H atoms | 2328 | 2162 | 8376 | 2060 | 1985 |
| Protein | 2069 | 1948 | 7810 | 1937 | 1918 |
| Ligand/ion | 5 | 36 | 47 | 6 | - |
| Water | 254 | 178 | 519 | 117 | 67 |
| B-factors (Å<sup>2</sup>) | 20.06 | 27.74 | 35.86 | 54.69 | 48.46 |
| Protein | 18.42 | 26.52 | 35.62 | 54.99 | 48.61 |
| Ligand/ion | 17.25 | 40.08 | 47.68 | 49.02 | - |
| Water | 33.45 | 38.61 | 38.35 | 49.97 | 44.27 |
| R.m.s. deviations |  |  |  |  |  |
| Bond lengths (Å) | 0.012 | 0.002 | 0.007 | 0.008 | 0.009 |
| Bond angles (°) | 1.25 | 0.65 | 0.87 | 1.07 | 1.06 |
| Ramachandran favored (%) | 97.52 | 96.64 | 96.79 | 95.3 | 95.3 |
| Ramachandran allowed (%) | 2.48 | 3.36 | 3.21 | 4.7 | 4.7 |
| Ramachandran outliers (%) | 0 | 0 | 0 | 0 | 0 |
| PDB ID | 9EZU | 9EZV | 9EZW | 9EZH | 9EZI |

<sup>a</sup> Each data set was collected from a single crystal. Values in parentheses correspond to the highest resolution shell. Values reported by XDS.988 <sup>b</sup> Values reported by Phenix

**Supplementary Table.8.**
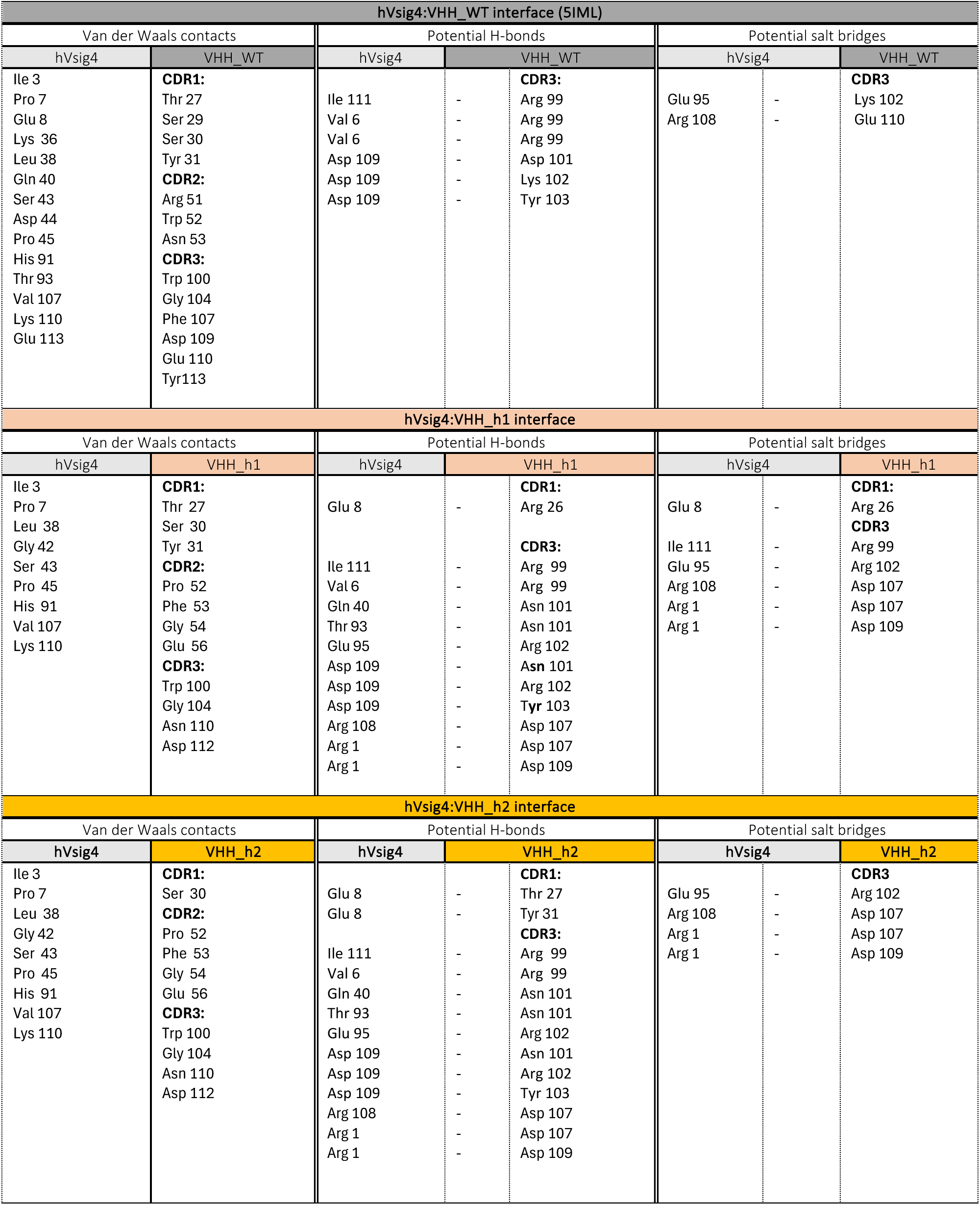
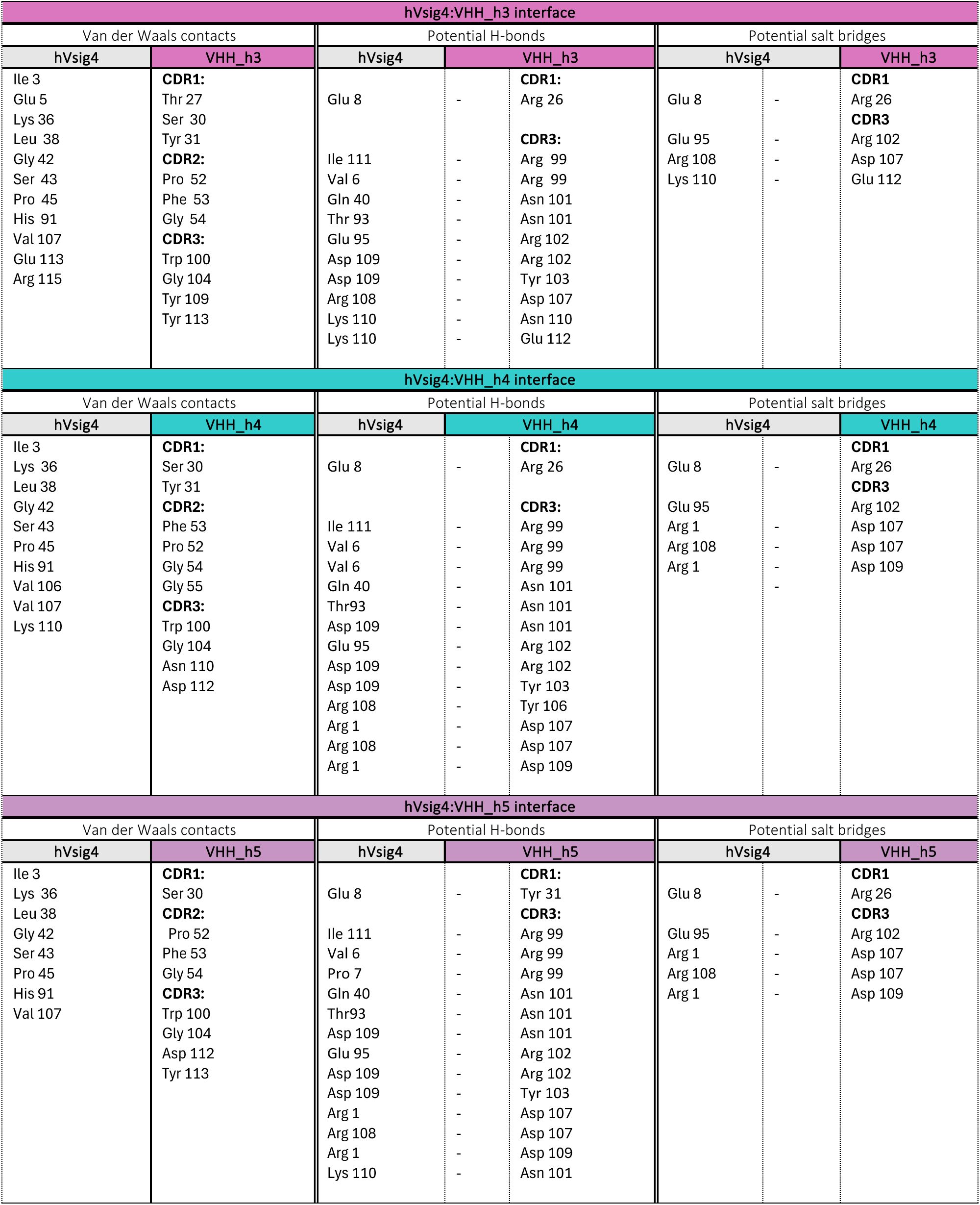
Interactions at the VHH_WT:hVSIG4 interfaces compared to the interactions at the VHH_h:hVSIG4 interfaces.

